# Identification of a neural basis orchestrating anorexia nervosa-like phenotypes

**DOI:** 10.64898/2026.02.07.704578

**Authors:** Cunjin Su, Yuanzhong Xu, Maojie Yang, Dan Liu, Yuhan Cao, Runzhou Yang, Caroline A. Regnauld, Emma C. Wheeler, Benjamin R. Arenkiel, Qingchun Tong

## Abstract

Anorexia nervosa (AN) is a severe, often fatal, psychiatric disorder characterized by self-imposed food restriction, hyperactivity, and profound affective disturbances. Despite its high morbidity and mortality, the neural mechanisms underlying AN remain poorly defined, largely because no animal model has fully recapitulated its core behavioral manifestations. Here, we demonstrate that aberrant activation of glutamatergic neurons in the mediobasal hypothalamus (MBH) is sufficient to induce AN-like phenotypes in mice, including self-starvation leading to death, hyperactivity, anhedonia, social avoidance, and heightened anxiety, which were reproduced by targeted activation of MBH neurons expressing steroidogenic factor 1 (SF1)/ estrogen receptor α (ERα). Mechanistically, the AN-like phenotypes were rescued by ablating release of glutamate or brain-derived neurotrophic factor (BDNF). Conversely, BDNF overexpression in SF1/ERα neurons was sufficient to promote AN-like behaviors through increasing glutamate release. Thus, BDNF-dependent enhancement of glutamatergic signaling from SF1/ERα neurons drives AN-associated phenotypes, providing a potential target for therapeutic intervention.

## Introduction

Normal body weight homeostasis is maintained by balanced food intake and energy expenditure, processes that are coordinated by a distributed neural network underlying feeding behavior and metabolism^1^. Disruption of these circuits can lead to profound abnormalities in feeding and metabolism. Anorexia nervosa (AN) is a severe psychiatric disorder characterized by persistent food restriction, extreme weight loss, one of the highest mortality rates among psychiatric illnesses, and commonly occurring during adolescents and females ^2^. In addition to voluntary restrictive feeding, AN is associated with a constellation of behavioral abnormalities, including anxiety, hyperactivity, social withdrawal, and anhedonia ^3–6 7–11^. Furthermore, stressful and/or traumatic early life events are widely recognized as important risk factors contributing to the onset of AN^4,12,13^. Accordingly, AN is generally viewed as a psychiatric disorder with profound metabolic and endocrine consequences^14^. Despite extensive clinical characterization, the neural mechanism underlying of AN remains poorly understood.

Studies on AN patients have identified numerous genetic, structural, behavioral, and physiological alterations associated with disease development ^2,15,16^. Among the implicated molecular pathways, brain-derived neurotrophic factor (BNDF) has emerged as a particularly potential candidate ^17^. BDNF is a member of the neurotrophic family that plays essential roles in neuronal development and synaptic plasticity ^18^. In addition, BDNF is a critical regulator of energy balance, and its expression within multiple brain sites contributes to body weight regulation ^19,20^. Human genetic studies have identified significant associations between AN and BDNF polymorphisms, particularly the BDNF^Val66Met^ variant ^21–23^. Consistently, animal models carrying this mutation exhibit heightened susceptibility to AN-like phenotypes under specific challenges ^21^. However, the contribution of this variant to AN development remains controversial because studies examining BDNF expression in AN patients have yielded inconsistent findings ^17,24^. Consequently, whether and how BNDF contributes to AN pathogenesis is still unclear.

Progress toward understanding the neural basis of AN has been further limited by the lack of animal models that faithfully recapitulate the core features of the disorder^12,25^. Most preclinical studies have relied on activity-based anorexia (ABA) paradigms, whereas others have employed genetic models involving BDNF variants ^21,26^. Although these models reproduce selected aspects of AN, none fully capture the defining clinical feature of voluntary restrictive feeding and self-starvation despite severe nutritional depletion^12^. As a result, the mechanisms driving AN-like behavior remain difficult to investigate experimentally.

The mediobasal hypothalamus (MBH) is a critical regulator of feeding behavior and energy balance. Within the MBH, glutamatergic neurons are highly enriched in the ventromedial hypothalamus (VMH) and are also present in neighboring hypothalamic regions. The MBH expresses numerous neuropeptides involved in body weight regulation, including BDNF^27,28^. The VMH contains several genetically defined neuronal populations, including steroidogenic factor-1 (SF1)- and estrogen receptor α (ERα)-expressing neurons, which partially overlap anatomically. These neuronal populations regulate diverse physiological and behavioral processes, including feeding, stress responses, locomotion, and social behaviors^29–32^, positioning them as compelling candidates for mediating AN-related phenotypes. Notably, glutamatergic ERα neurons are also distributed throughout neighboring MBH regions.

Here, we demonstrate that chronic activation of MBH glutamatergic neurons or VMH SF1/ERα neurons induces voluntary self-starvation, hyperactivity, anxiety, social avoidance, and anhedonia, collectively recapitulating the core behavioral features of AN. Mechanistically, our findings reveal that SF1 and ERα neurons both contribute to restrictive feeding, anxiety, and anhedonia, whereas ERα neurons are specifically required for hyperactivity. Furthermore, we identify enhanced BDNF signaling and consequent increases in glutamatergic transmission from VMH SF1/ERα neurons as key drivers of AN-like phenotypes. These findings establish a robust animal model that reproduces the major behavioral manifestations of AN and provides mechanistic insight into the neural circuitry underlying disease pathogenesis.

## Results

### *Chronic activation of* MBH glutamatergic neurons induces AN-like phenotypes

Neurological and psychiatric disorders are perceived to arise from persistent alterations in neuronal activity states^33^. We previously adopted a strategy for chronically activating genetically defined neuronal populations through cell type-specific expression of the bacterial voltage-gated sodium channel NaChBac^34–37^. To examine the consequences of sustained activation of MBH glutamatergic neurons, we delivered Cre-dependent AAV-DIO-NaChBac-GFP or control vectors into the MBH of *Vglut2-Cre* mice. To validate neuronal activation, AAV-DIO-NaChBac-GFP was injected unilaterally into the MBH, while a control AAV-DIO-mCherry virus was injected into the contralateral hemisphere (Fig. 1a). NaChBac expression resulted in a marked increase in c-Fos immunoreactivity relative to the control side, indicating elevated neuronal activity (Fig. 1b). Bilateral MBH delivery of NaChBac similarly produced robust c-Fos induction compared with AAV-DIO-EYFP controls (Fig. 1c; Supplementary Fig. S1a), confirming effective chronic activation of MBH glutamatergic neurons.

**Figure 1.**
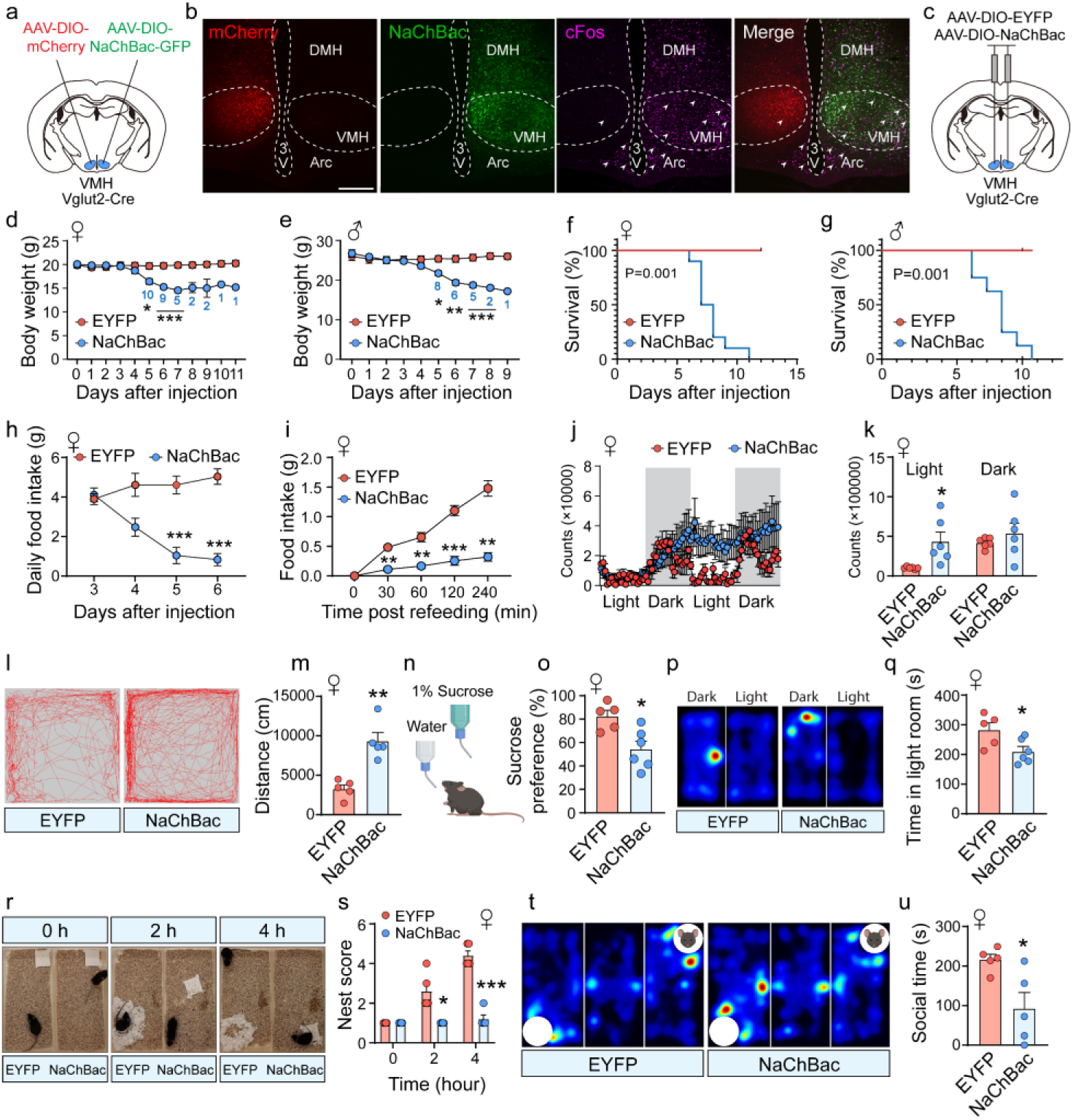
Chronic activation of VH glutamatergic neurons induces anorexia-like phenotypes. **(a)** Schematic of unilateral viral delivery (AAV-DIO-mCherry, or AAV-DIO-NaChBac) into the VH of Vglut2-Cre mice. **(b)** Representative images from unilateral injections in the same brain section showing expression of mCherry, NaChBac-GFP, and robust c-Fos induction (arrows) on the NaChBac-injected side. Scale bar, 300 μm. **(c)** Schematic of bilateral viral delivery (AAV-DIO-EYFP, and AAV-DIO-NaChBac) into the VH of Vglut2-Cre mice. **(d, e)** Body weight trajectories of female **(d)** and male **(e)** Vglut2-Cre mice after bilateral VH injection of AAV-DIO-NaChBac or AAV-DIO-GFP. Females, n = 10 per group; males, n = 8 per group; two-way ANOVA; *P < 0.05, **P < 0.01, ***P < 0.001. **(f, g)** Kaplan–Meier survival analysis of female **(f)** and male **(g)** mice expressing GFP or NaChBac. Mice were considered as non-survial when body weight loss reached 30%, log-rank test, P < 0.001 for both sexes. **(h)** Daily food intake in EYFP- and NaChBac-expressing female mice. n = 6 per group; two-way ANOVA; ***P < 0.001. **(i)** Cumulative food intake during fasting–refeeding in EYFP- and NaChBac-expressing female mice. n = 5 per group; two-way ANOVA; **P < 0.01, ***P < 0.001. **(j)** Real-time analysis of spontaneous locomotor activity of female mice measured in metabolic cages (TSE). **(k)** Quantification of locomotor activity during light and dark phases. n = 6 per group; unpaired two-tailed t test; P = 0.017 (light), P = 0.387 (dark). **(l, m)** Open-field test. **(l)** Representative movement trajectories. **(m)** Total distance traveled in 15 min. n = 5 per group; unpaired two-tailed t test, P=0.001. **(n)** Schematic of the sucrose preference test. **(o)** Sucrose preference in EYFP- and NaChBac-expressing female mice. n = 5 (EYFP), n = 6 (NaChBac); unpaired two-tailed t test, P=0.011. **(p, q)** Light–dark box test. Representative trajectories **(p)**, time spent in the light room **(q)**. n = 5 (EYFP), n = 6 (NaChBac); unpaired two-tailed t test, P=0.031. **(r,s)** Nesting behavior representative images **(r)**, nesting scores over time **(s)** . n = 5 per group; two-way ANOVA, P=0.048 at 2 h, P<0.001 at 4 h. **(t, u)** Three-chamber sociability test. Representative trajectories **(t)**, time spent interacting with a stranger mouse **(u)**. n = 5 per group; unpaired two-tailed t test; P=0.019.

We next examined the physiological and behavioral consequences of this manipulation. Chronic activation of MBH glutamatergic neurons produced a rapid and progressive loss of body weight beginning approximately five days after viral delivery in both female (Fig. 1d) and male mice (Fig. 1e), ultimately resulting in mortality (Figs. 1f and 1g). This phenotype was associated with a pronounced voluntary reduction in food intake, as measured by ad libitum consumption and fasting-induced refeeding assays (Figs. 1h and 1i). Importantly, reduced food intake was not secondary to impaired locomotor function. On the contrary, NaChBac-expressing females exhibited increased locomotor activity, reflected by elevated beam-break counts in metabolic monitoring chambers (Figs. 1j and 1k) and increased movement in the open-field test (OFT, Figs. 1l and 1m). These findings resemble the paradoxical hyperactivity frequently observed in AN patients.

To determine whether chronic activation of MBH glutamatergic neurons also reproduces affective and social abnormalities characteristic of AN, we assessed anhedonia, anxiety-related behavior, and sociability. In the sucrose preference test, NaChBac-expressing female mice exhibited a marked reduction in sucrose consumption relative to controls (Figs. 1n and 1o), indicative of anhedonia. In the light-dark test (LDT), these mice spent significantly less time in the illuminated compartment (Figs. 1p and 1q), suggesting heightened anxiety-like behavior. Similarly, nest-building performance was significantly impaired (Figs. 1r and 1s), providing additional evidence of increased anxiety. We next evaluated sociability using a three-chamber social interaction assay. Compared with control animals, NaChBac-expressing mice spent significantly less time interacting with a novel conspecific (Figs. 1t and 1u), indicating impaired sociability and increased social avoidance.

Collectively, these findings demonstrate that chronic activation of MBH glutamatergic neurons induces lethal voluntary self-starvation, hyperactivity, anxiety-like behavior, social avoidance, and anhedonia, closely recapitulating the core features of AN. Notably, although NaChBac-expressing mice displayed no abnormal jumping behavior in their home cages (**Supplementary Video 1**), exposure to a novel environment elicited frequent jumping episodes (**Supplementary Video 2**), suggesting heightened stress sensitivity and anxiety. This observation is particularly relevant given the established association between stressful life events and the onset of AN in humans^21,38^. Importantly, comparable phenotypes were observed in male mice (Supplementary Figs. S1b-S1n), indicating that these effects are not restricted to females.

### Chronic activation of MBH SF1 and ERa neurons elicits typical AN-like phenotypes

Because stereotaxic targeting of the MBH in *Vglut2-Cre* mice likely encompasses glutamatergic neurons outside the VMH, we next sought to determine whether more precisely defined VMH neuronal populations contribute to the observed phenotypes. SF1 expression provides a specific molecular marker for VMH neurons, while ERα expression partially overlaps with SF1 within the VMH, particularly in the ventrolateral subdivision, and extends into adjacent MBH regions^39^. Consistent with previous reports, female mice possessed a greater number of ERα-expressing neurons than males (Supplementary Fig. S2a). To selectively target these populations, we generated compound *SF1-Cre::ERα-Cre* mice and delivered AAV-DIO-NaChBac-GFP into the VMH. Mice expressing either SF1-Cre or ERα-Cre alone served as comparison groups (Fig. 2a). Robust NaChBac expression and c-Fos induction were observed in both compound and single-driver lines (Fig. 2b). Consistent with the partial overlap between SF1 and ERα populations, compound mice exhibited broader NaChBac and c-Fos expression throughout the VMH and adjacent MBH regions relative to either single-driver line (Fig. 2b), confirming successful chronic activation of the combined SF1/ERα neuronal population.

**Figure. 2.**
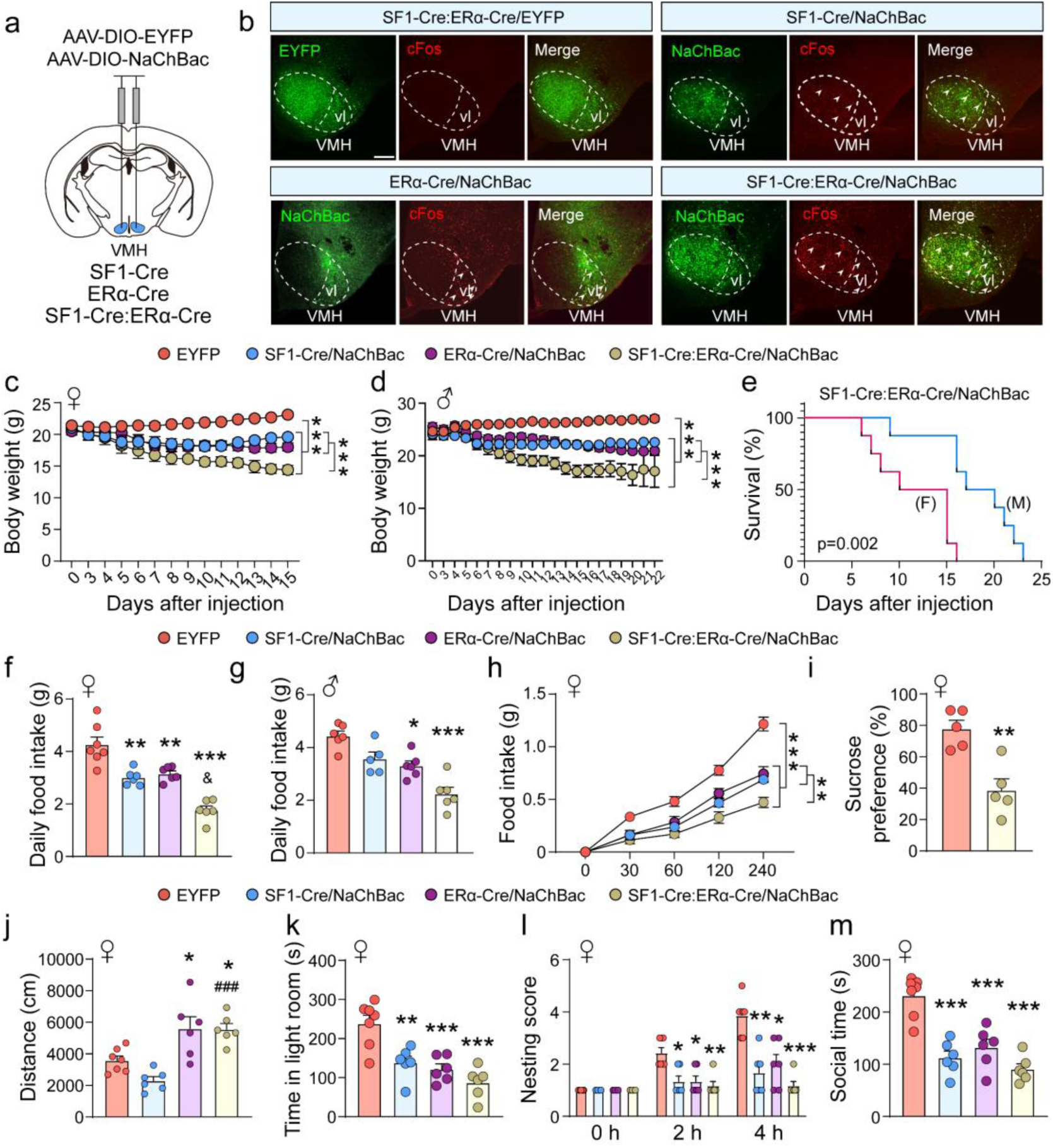
Chronic activation of VMH SF1 and ERa neurons leads to typical anorexia like behaviors. **(a)** Schematic of viral delivery (AAV-DIO-EYFP or AAV-DIO-NaChBac) into the VMH of SF1-Cre, ERa-Cre, and SF1-Cre::ERa-Cre mice. **(b)** Representative images showing EYFP or NaChBac expression patterns in VMH, and c-Fos induction (arrows) in NaChBac-expressing mice. Scale bar, 250 μm. **(c, d)** Body weight trajectories of female **(c)** and male **(d)** SF1-Cre, ERa-Cre, and SF1-Cre::ERa-Cre mice after VMH injection of AAV-DIO-NaChBac or AAV-DIO-EYFP. Females, n=8 (EYFP), n=6 (SF1-Cre/NaChBac), n=6 (ERa-Cre/NaChBac), n=8 (SF1-Cre::ERa-Cre/NaChBac); males, n=6 (EYFP), n=6 (SF1-Cre/NaChBac), n=8 (ERa-Cre/NaChBac), n=8 (SF1-Cre::ERa-Cre/NaChBac); two-way ANOVA, ***P<0.001. **(e)** Kaplan–Meier survival analysis of female and male SF1-Cre::ERa-Cre mice expressing NaChBac. Mice were considered as non-survival when body weight loss reached 30%, log-rank test, P=0.021 between males and females. **(f, g)** Daily food intake in females **(f)** and males **(g)** EYFP- and NaChBac-expressing mice. Females, n=7 (EYFP), n=6 (SF1-Cre/NaChBac), n=6 (ERa-Cre/NaChBac), n=6 (SF1-Cre::ERa-Cre/NaChBac); males, n=6 (EYFP), n=5 (SF1-Cre/NaChBac), n=6 (ERa-Cre/NaChBac), n=6 (SF1-Cre::ERa-Cre/NaChBac); two-way ANOVA; *P<0.05, **P<0.01, ***P<0.001 versus EYFP group; ^&^ P=0.023 versus ERa-Cre/NaChBac group. **(h)** Cumulative food intake during fasting–refeeding in EYFP- and NaChBac-expressing female mice. n=5 (EYFP), n=5 (SF1-Cre/NaChBac), n=7 (ERa-Cre/NaChBac), n=6 (SF1-Cre::ERa-Cre/NaChBac); two-way ANOVA, **P=0.007, ***P<0.001. **(i)** Sucrose preference in EYFP- and NaChBac-expressing female SF1-Cre::ERa-Cre mice. n = 5 per group; unpaired t test, P=0.003. **(j-m)** Total distance traveled in open the field test **(j)**, time spent in the light room in light-dark test **(k)**, nesting score **(l)**, and social time with stranger in three-chamber test **(m)** in female mice. n=7 (EYFP), n=6 (SF1-Cre/NaChBac), n=6 (ERa-Cre/NaChBac), n=6 (SF1-Cre::ERa- Cre/NaChBac); two-way ANOVA; *P<0.05, **P<0.01, ***P<0.001 versus EYFP; ^###^P<0.001 versus SF1-Cre/NaChBac.

Chronic activation of SF1/ERα neurons produced a progressive reduction in body weight that ultimately resulted in mortality in both females (Figs. 2c and 2e) and males (Figs. 2d and 2e). In contrast, activation of SF1 or ERα neurons alone resulted in more modest body weight reductions without lethality (Figs. 2c and 2d). Notably, female mice consistently succumbed earlier than males (Fig. 2e), suggesting a sexually dimorphic effect that may reflect the greater abundance of ERα-expressing neurons in females. The profound weight loss observed in SF1/ERα mice was associated with markedly reduced food intake in both females (Fig. 2f) and males (Fig. 2g), and diminished fasting-induced refeeding responses (Fig. 2h). Consistent with the body weight phenotype, reductions in food intake were substantially greater in double-driver mice than in either single-driver group (Figs. 2f-2h).

Behavioral analyses further revealed that activation of SF1/ERα neurons reduced sucrose preference (Fig. 2i), indicative of anhedonia. Interestingly, activation of SF1 neurons alone decreased locomotor activity, whereas activation of ERα neurons alone or the combined SF1/ERα population increased locomotion in the open-field test (Fig. 2j), suggesting a specific role for ERα neurons in promoting locomotor activity. Despite increased locomotion, SF1/ERα activation produced a pronounced reduction in time spent in the illuminated compartment during the LDT (Fig. 2k), consistent with heightened anxiety-like behavior. Similarly, mice with SF1/ERα activation exhibited marked deficits in nest building (Fig. 2l) and sociability (Fig. 2m), whereas activation of either SF1 or ERα neurons alone produced comparatively modest effects. Equivalent behavioral changes were also observed in male mice (Supplementary Figs. S2b-S2e). Together, these findings demonstrate that chronic activation of SF1/ERα neurons is sufficient to reproduce the core behavioral features of AN. While SF1 and ERα neuronal populations each contribute to restrictive feeding, anxiety-like behavior, and anhedonia, ERα neurons appear to play a unique role in mediating hyperactivity, a hallmark feature of AN.

### Glutamate and BDNF are both required for AN-like phenotypes

VMH neurons release multiple signaling molecules, including the fast excitatory neurotransmitter glutamate and the neurotrophin BDNF^40^. To determine whether glutamatergic transmission is required for the AN-like phenotypes elicited by SF1/ERα neuron activation, we used a CRISPR-Cas9 strategy to selectively disrupt Vglut2 by delivering either AAV-DIO-Cas9-sgVglut2^41^ or a control AAV-DIO-Cas9-sgRNA vector into the VMH of *SF1-Cre::ERα-Cre* mice. To verify the effectiveness of this manipulation, we monitored optogenetically evoked glutamate release using fiber photometry and the glutamate sensor GluSnFR3.0 expressed in the lateral septum (LS), a major downstream target of MBH neurons (Fig. 3a). Expression of the red-light-activated opsin ChrimsonR in SF1/ERα projections through delivery of AAV-DIO-ChrimsonR viral vectors of the VMH and GluSnFR3.0 in LS neurons through delivery of AAV-Cre-GFP and AAV-DIO-GluSnFR3.0 viral vectors to the LS was confirmed histologically (Fig. 3b). Whereas red-light stimulation produced robust glutamate sensor responses in control mice, mice receiving sgVglut2 showed virtually no detectable response (Fig. 3c-3e), indicating near-complete elimination of glutamate release following Cas9-mediated deletion of Vglut2.

**Figure. 3.**
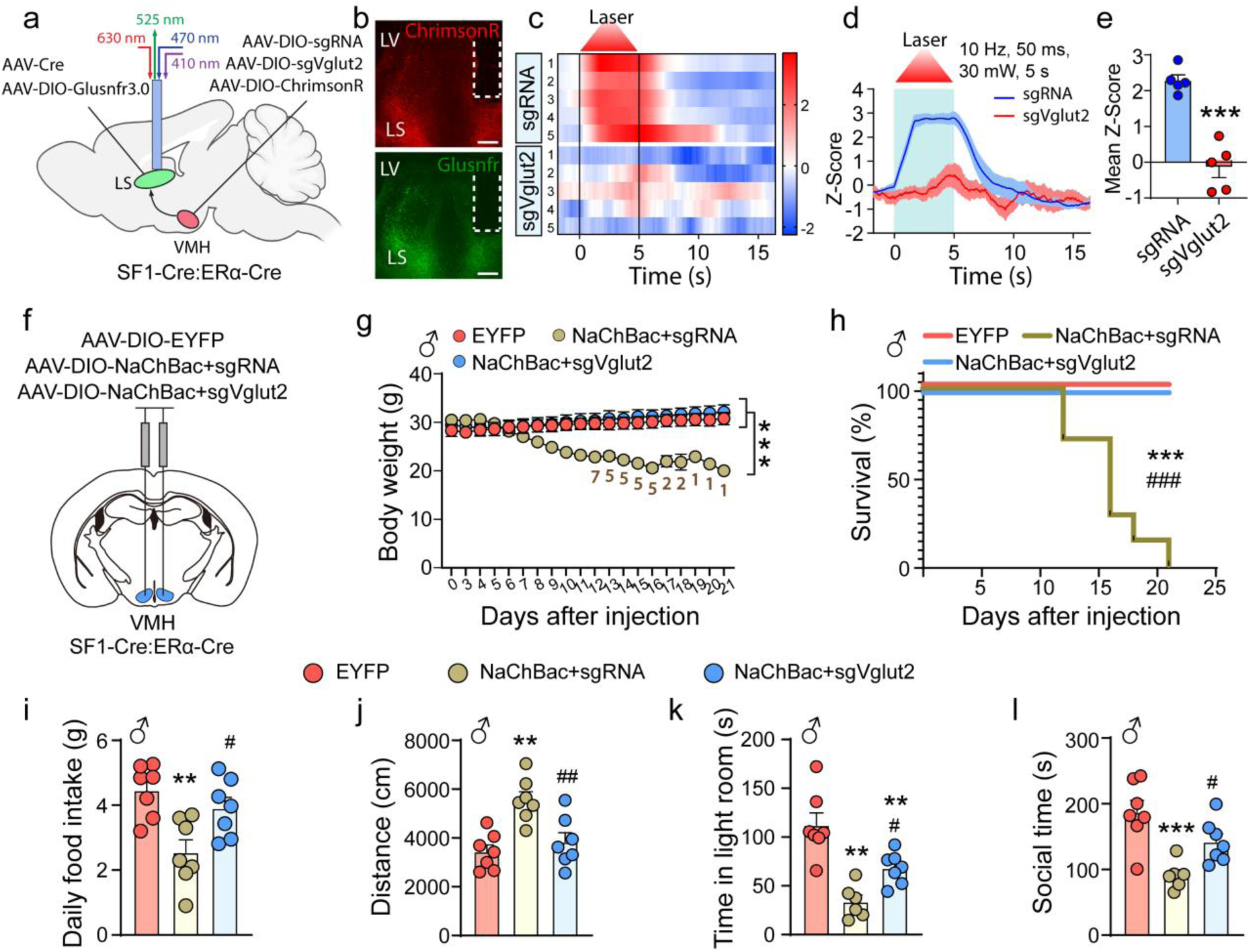
Glutamate release is required for anorexia phenotypes. **(a)** Diagram of in vivo optogenetic stimulation of VMH SF1 and ERα neuronal terminals with simultaneous fiber photometry recording of glutamate sensor signals in downstream lateral septum neurons.**(b)** Representative Glusnfr (green) and ChRimsonR (terminals from VMH, red) expression in lateral septum. Scale bar, 250 μm. **(c)** Heatmap of glutamate release Z score signals of individual trails in response to 5-s photostimulation. **(d)** Averaged trace of glutamate release Z score signals of lateral septum neurons before, during, and after 5-s stimulation of SF1 and ERa neurons axons in VMH. **(e)** Mean of glutamate release Z score during photostimulation. n=5 per group, unpaired two-tailed t test, P<0.001. **(f)** Diagram of viral delivery (AAV-DIO-EYFP, AAV-DIO-NaChBac + AAV-DIO-sgRNA, AAV-DIO-NaChBac + AAV-DIO-sgVglut2) into the VMH of SF1-Cre::ERa-Cre male mice. **(g)** Body weight trajectories of male SF1-Cre::ERa-Cre mice. n=7 per group, two-way ANOVA, ***P<0.001. Numbers shown in the figure represent the number of remaining surviving mice included in the analysis at each time point. **(h)** Kaplan–Meier survival analysis of mice in EYFP, NaChBac+sgRNA, and NaChBac+sgVglut2 groups. Mice were considered non-survival when body weight loss reached 30%, log-rank test, ***P<0.001 versus EYFP, ^###^P<0.001 versus NaChBac+sgVglut2. **(i)** Daily food intake. n=7 per group, one-way ANOVA, **P=0.003 versus EYFP, ^#^P=0.031 versus NahBac+sgRNA. **(j-l)** Total distance traveled in open field test **(j)**, time spent in the light room in light-dark test **(k)**, social time with stranger in three-chamber test **(l).** n=7 per group, one-way ANOVA, **P<0.01, ***P<0.001 versus EYFP; ^#^P<0.05, ^##^P<0.01 versus NaChBac+sgRNA.

Having confirmed efficient suppression of glutamatergic transmission, we next examined its functional role in NaChBac-induced AN-like phenotypes by co-injecting AAV-DIO-NaChBac-GFP with either control sgRNA or sgVglut2 into the VMH (Fig. 3f). As expected, mice receiving NaChBac together with control sgRNA displayed progressive body weight loss beginning approximately five days after viral delivery (Fig. 3g). Strikingly, deletion of Vglut2 completely rescued both the severe weight loss and lethality induced by chronic activation of SF1/ERα neurons (Figs. 3g and 3h). In addition, loss of glutamate release reversed voluntary food restriction (Fig. 3i), hyperactivity (Fig. 3j), anxiety-like behavior in LDT (Fig. 3k), and reduced sociability (Fig. 3l). Together, these findings demonstrate that enhanced glutamatergic output is required for the AN-like phenotypes produced by chronic activation of SF1/ERα neurons.

We next investigated the contribution of BDNF signaling. To this end, we generated *SF1-Cre::ERα-Cre::BDNF^flox/flox^* mice and delivered AAV-DIO-NaChBac-GFP into the VMH (Fig. 4a).

**Figure. 4.**
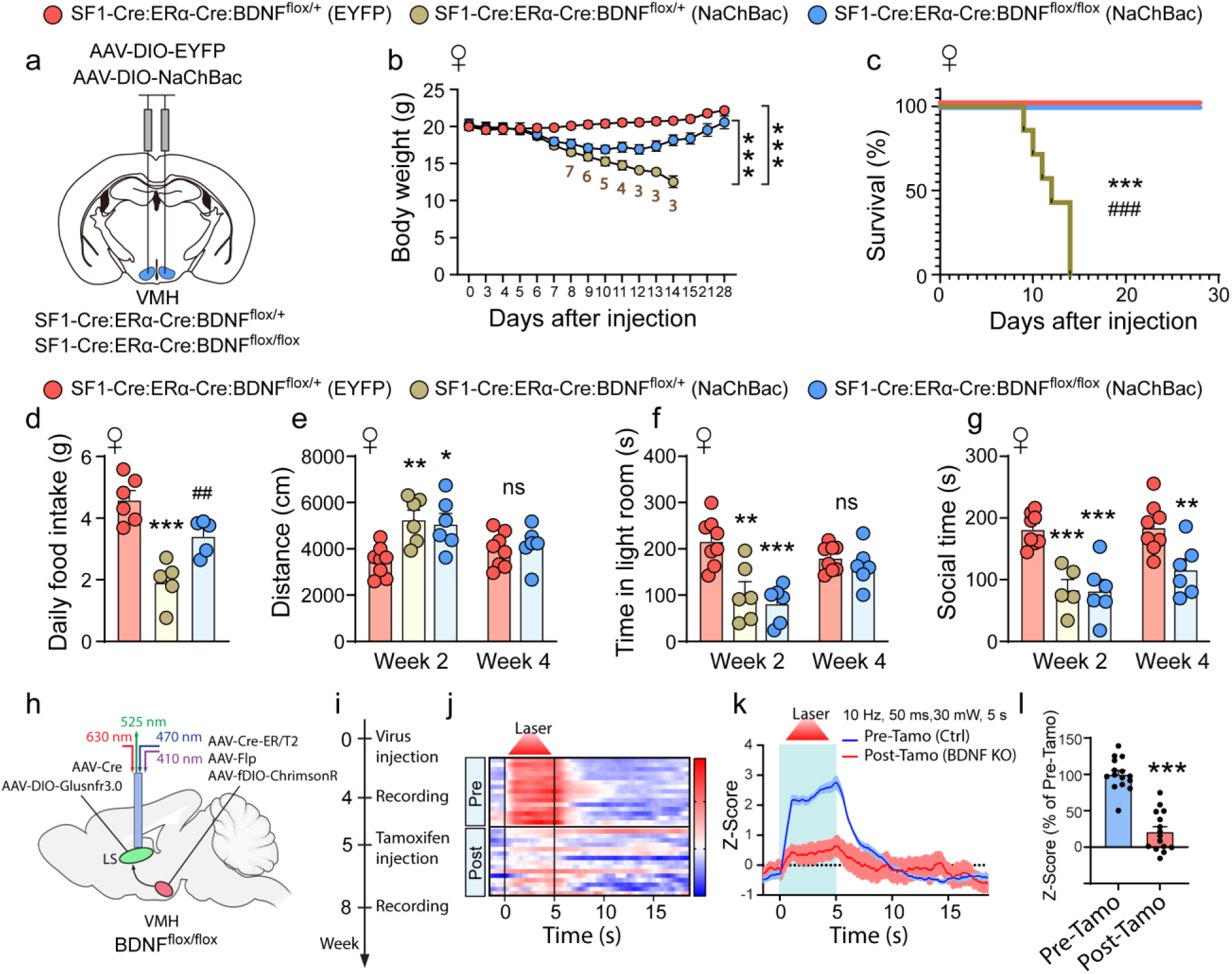
BDNF release is required for anorexia phenotypes. **(a)** Diagram of viral delivery (AAV-DIO-EYFP, AAV-DIO-NaChBac) into VH of SF1-Cre::ERa-Cre::BDNFflox/+and SF1-Cre::ERa-Cre::BDNFflox/flox female mice. **(b)** Body weight trajectories of female mice. n=8 (SF1-Cre::ERa-Cre::BDNFflox/++EYFP), n=6 (SF1-Cre::ERa-Cre::BDNFflox/++NaChBac), n=6 ( SF1-Cre::ERa-Cre::BDNFflox/flox+NaChBac ); two-way ANOVA, ***P<0.001. Numbers shown in the figure represent the number of mice included in the analysis at each time point. **(c)** Kaplan–Meier survival analysis of mice in SF1-Cre::ERa-Cre::BDNFflox/++EYFP, SF1-Cre::ERa-Cre::BDNFflox/++NaChBac, SF1-Cre::ERa-Cre::BDNFflox/flox+NaChBac groups. Mice were considered as non-survival when body weight loss reached 30%, log-rank test, ***P<0.001 versus EYFP, ###P<0.001 versus SF1-Cre::ERa-Cre::BDNFflox/flox+NaChBac. **(d)** Daily food intake. n=5-6 per group, one-way ANOVA, ***P<0.001 versus EYF; ##P=0.009 versus SF1-Cre::ERa-Cre::BDNFflox/++NaChBac. **(e-g)** Total distance traveled in open field test **(e)**, time spent in the light room in light-dark test **(f)**, social time with stranger in three-chamber test **(g)**. n=6-8 per group, one-way ANOVA, *P<0.05, **P<0.01, ***P<0.001 versus EYFP. **(h)** Diagram of in vivo optogenetic stimulation of VMH SF1 and ERα neuronal terminals with simultaneous fiber photometry recording of glutamate sensor signals in downstream lateral septum neurons before and after inducible BDNF knockout. **(i)** Schematic timeline of fiber photometry recording and tamoxifen injection. **(j)** Heatmap of glutamate release Z score signals of individual trails in response to 5-s optogenetic stimulation. **(k)** Averaged trace of glutamate release Z score signals of lateral septum neurons before, during, and after 5-s stimulation of SF1 and ERa neurons axons in VMH before (blue line) and after inducible BDNF knockout (red line). **(l)** Mean of glutamate release Z score during optogenetic stimulation. n=15 trails per group, unpaired two-tailed t test, P<0.001.

Efficient deletion of BDNF was confirmed in the VMH of *SF1-Cre::ERα-Cre::BDNF^flox/flox^*mice relative to *SF1-Cre::ERα-Cre::BDNF^flox/+^* controls (Supplementary Fig. S3). Notably, BDNF deletion did not alter baseline body weight during the age range examined in this study (Fig. 4b). Consistent with our previous findings, NaChBac expression in *SF1-Cre::ERα-Cre::BDNF^flox/+^* mice induced progressive body weight loss beginning approximately five days after viral delivery and ultimately resulted in mortality (Figs. 4b,c). In contrast, *SF1-Cre::ERα-Cre::BDNF^flox/flox^*mice exhibited only a modest initial reduction in body weight, followed by gradual recovery, achieving near-complete normalization by 28 days after viral delivery (Fig. 4b). Correspondingly, no lethality was observed in mice lacking BDNF (Fig. 4c), indicating that BDNF deletion markedly attenuates the pathological consequences of chronic SF1/ERα neuron activation.

Consistent with these effects on body weight, BDNF deletion rescued NaChBac-induced voluntary food restriction (Fig. 4d). It also reversed hyperactivity (Fig. 4e) and anxiety-like behavior in LDT (Fig. 4f), although these effects emerged only four weeks after viral delivery and were not evident at earlier time points. In contrast, deficits in sociability were only partially improved and were not fully rescued at any time examined (Fig. 4g).

The similar rescue produced by either BDNF deletion or blockade of glutamate release raised the possibility that BDNF acts upstream of glutamatergic transmission. To test this hypothesis directly, we assessed glutamate release following inducible deletion of BDNF. We delivered Tam (tamoxifen)-inducible AAV-Cre-ER together with AAV-Flp and AAV-fDIO-ChrimsonR to the MBH of *BDNF^flox/flox^* mice and injected AAV-Cre-GFP plus AAV-DIO-GluSnFR3.0 into the LS (Fig. 4h). This strategy enabled tamoxifen-dependent deletion of BDNF while simultaneously permitting optogenetic stimulation of MBH neurons and monitoring of glutamate release in the LS. Glutamate sensor activity was recorded both before and after tamoxifen administration within the same animals (Fig. 4i). Notably, tamoxifen-induced deletion of BDNF markedly reduced optogenetically evoked glutamate release, as demonstrated by population heat maps (Fig. 4j), representative recordings (Fig. 4k), and quantitative analysis (Fig. 4l). Collectively, these results demonstrate that both glutamate and BDNF signaling are required for the AN-like phenotypes induced by chronic activation of SF1/ERα neurons. Moreover, the reduction in glutamate release following BDNF deletion supports a model in which enhanced BDNF signaling promotes AN-like behaviors, at least in part, by potentiating glutamatergic transmission from SF1/ERα neurons.

### Augmented BDNF action from MBH glutamatergic neurons promotes AN-like phenotypes

Given that BDNF deletion attenuated both AN-like phenotypes and glutamate release, we next investigated whether enhanced BDNF signaling contributes directly to the pathological effects of chronic MBH glutamatergic neuron activation. To address this question, we injected AAV-DIO-NaChBac-GFP into one side of the MBH and control AAV-DIO-mCherry into the contralateral side of *Vglut2-Cre* mice (Fig. 5a). Immunohistochemical analysis revealed significantly higher BDNF expression on the NaChBac-injected side compared with the control side (Figs. 5b and 5c), suggesting that chronic activation of MBH glutamatergic neurons upregulates BDNF expression and implicating enhanced BDNF signaling in the development of AN-like phenotypes.

**Figure. 5.**
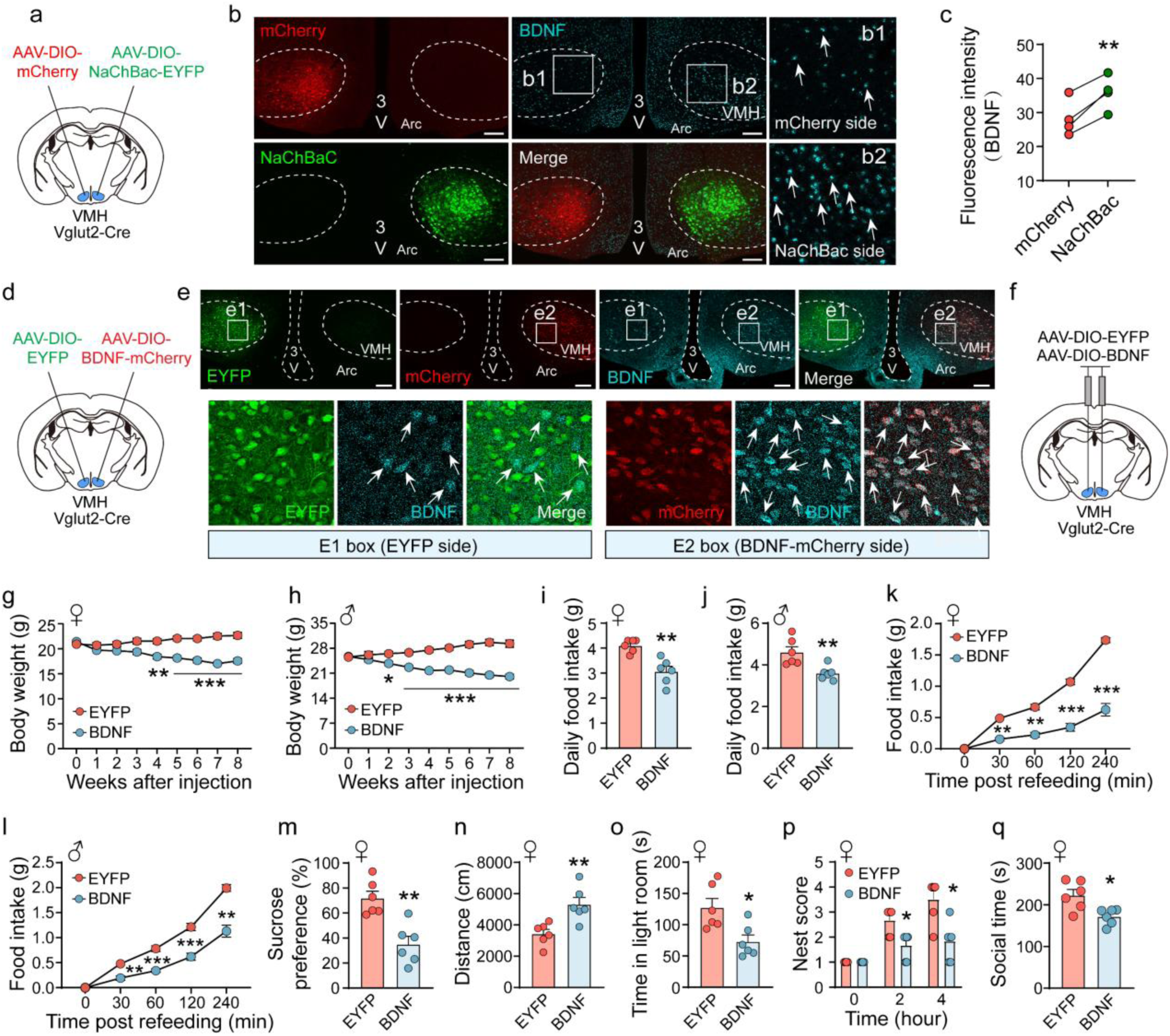
Augmented BDNF action from VMH glutamatergic neurons promoted anorexia like behaviors. **(a)** Diagram of unilateral viral delivery (AAV-DIO-mCherry, AAV-DIO-NaChBac-EYFP) into the VMH of Vglut2-Cre mice. **(b)** Representative images from unilateral injections in the same brain section showing mCherry (red), NaChBac-EYFP (green) expression, and BDNF immunostaining (cyan). **(b1)** a higher-magnification view of the left boxed area on mCherry injection side. **(b2)** a higher-magnification view of the right boxed area on NaChBac-EYFP injection side. Arrows indicate BDNF immunostaining positive cells. Scale bar, 250 μm. **(c)** Fluorescence intensity of BDNF in VMH. n=4, paired two-tailed t test, P<0.001. **(d)** Diagram of unilateral viral delivery (AAV-DIO-EYFP, AAV-DIO-BDNF-mCherry) into the VMH of Vglut2-Cre mice. **(e)** Representative images from unilateral injections in the same brain section showing EYFP (green), BDNF-mCherry (red) expression, and BDNF immunostaining (cyan). **(e1)** a higher-magnification view of the left boxed area on the AAV-DIO-EYFP injection side. **(e2)** a higher-magnification view of the right boxed area on the AAV-DIO-BDNF-mCherry injection side. Arrows indicate BDNF immunostaining positive cells. Scale bar, 250 μm. **(f)** Diagram of bilateral viral delivery (AAV-DIO-EYFP, AAV-DIO-BDNF-mCherry) into the VMH of Vglut2-Cre mice. **(g, h)** Body weight trajectories of female **(g)** and male **(h)** Vglut2-Cre mice after VMH injection of AAV-DIO-EYFP or AAV-DIO-BDNF. Females, n=6 per group; males, n=10 (EYFP), n=9 (BDNF); two-way ANOVA, *P<0.05, **P<0.01, ***P<0.001. **(i, j)** Daily food intake in EYFP- and BDNF-expressing Vglut2-Cre female **(i)** or male **(j)** mice. n=6 per group for both sexes; unpaired t test; P=0.001 (female), P=0.006 (male). **(k, l)** Cumulative food intake during fasting–refeeding in EYFP- and BDNF-expressing female **(k)** or male **(l)** Vglut2-Cre mice. Females, n = 5 per group; males, n=6-7 per group; two-way ANOVA; **P < 0.01, ***P < 0.001. **(m)** Sucrose preference in EYFP- and BDNF- expressing female Vglut2-Cre mice. n=6 per group; unpaired two-tailed t test, P=0.001. **(n-q)** Total distance traveled in the open field test **(n)**, time spent in the light room in light-dark test **(o)**, nesting behavior **(p)**, social time with stranger in three-chamber test **(q)** in female mice. n=6 per group; unpaired two-tailed t test **(n, o, q)**, two-way ANOVA **(p)**; *P<0.05, **P<0.01 versus EYFP.

To determine whether increased BDNF signaling is sufficient to induce AN-like behaviors, we generated a Cre-dependent AAV-DIO-BDNF-p2A-mCherry vector for conditional BDNF overexpression. We first validated viral efficacy by injecting AAV-DIO-BDNF-p2A-mCherry into one side of the hypothalamus and a control GFP vector into the contralateral side (Fig. 5d). After two weeks, robust colocalization of BDNF and mCherry was observed on the BDNF-injected side, whereas only low endogenous BDNF expression was detected on the control side (Fig. 5e), confirming efficient viral-mediated BDNF overexpression.

We next bilaterally delivered the BDNF virus to the VMH of *Vglut2-Cre* mice (Fig. 5f) and verified accurate targeting of the hypothalamus (Supplementary Fig. S4a). BDNF-overexpressing mice exhibited a progressive decline in body weight, reaching approximately 20% weight loss by eight weeks after viral injection in both females (Fig. 5g) and males (Fig. 5h). This reduction was accompanied by a marked decrease in daily food intake (Figs. 5i and 5j) and impaired fasting-induced refeeding responses (Figs. 5k and 5l), indicating voluntary restrictive feeding. In addition to reduced food intake, female mice overexpressing BDNF displayed a significant decrease in sucrose preference (Fig. 5m), consistent with anhedonia, and increased locomotor activity in OFT (Fig. 5n). Despite their elevated locomotion, these mice spent significantly less time in the illuminated compartment of LDT (Fig. 5o) and exhibited impaired nest-building behavior (Fig. 5p), indicative of heightened anxiety-like behavior. Furthermore, BDNF-overexpressing mice displayed spontaneous and vigorous jumping behavior when exposed to a novel cage (**Supplementary Video 3**), a response absent in control animals. This phenotype suggests increased sensitivity to novel or stressful environments and an exaggerated anxiety-related response. Consistent with these findings, female mice overexpressing BDNF spent significantly less time interacting with a novel conspecific in the three-chamber social interaction assay (Fig. 5q), indicating reduced sociability and enhanced social avoidance. Collectively, these behavioral and physiological alterations demonstrate that increased BDNF signaling in MBH glutamatergic neurons is sufficient to induce a broad spectrum of AN-like phenotypes, including voluntary food restriction, weight loss, hyperactivity, anxiety-like behavior, social avoidance, and anhedonia. Notably, equivalent phenotypes were observed in male mice (Supplementary Figs. S4b-S4j), suggesting that enhanced BDNF signaling engages a common neural mechanism capable of promoting AN-like behaviors in both sexes.

### Augmented BDNF action from SF1/ERa neurons promotes AN-like phenotypes

To determine whether enhanced BDNF signaling within SF1/ERα neurons is sufficient to induce AN-like phenotypes, we delivered AAV-DIO-BDNF-p2A-mCherry vectors into the VMH of *SF1-Cre::ERα-Cre* mice, as well as SF1-Cre and ERα-Cre mice, with AAV-DIO-GFP serving as a control (Fig. 6a). Successful and selective overexpression of BDNF was confirmed in SF1, ERα, and SF1/ERα neuronal populations (Fig. 6b). Targeted BDNF overexpression in SF1, ERα, or SF1/ERα neurons produced a progressive reduction in body weight in both female (Fig. 6c) and male mice (Fig. 6d). Notably, the magnitude of weight loss was greatest in SF1/ERα mice, indicating an additive contribution of these overlapping neuronal populations. Consistent with the body weight phenotype, BDNF overexpression significantly reduced daily food intake and fasting-induced refeeding responses in both sexes (Figs. 6e and 6h), demonstrating a robust voluntary restrictive feeding effect.

**Figure. 6.**
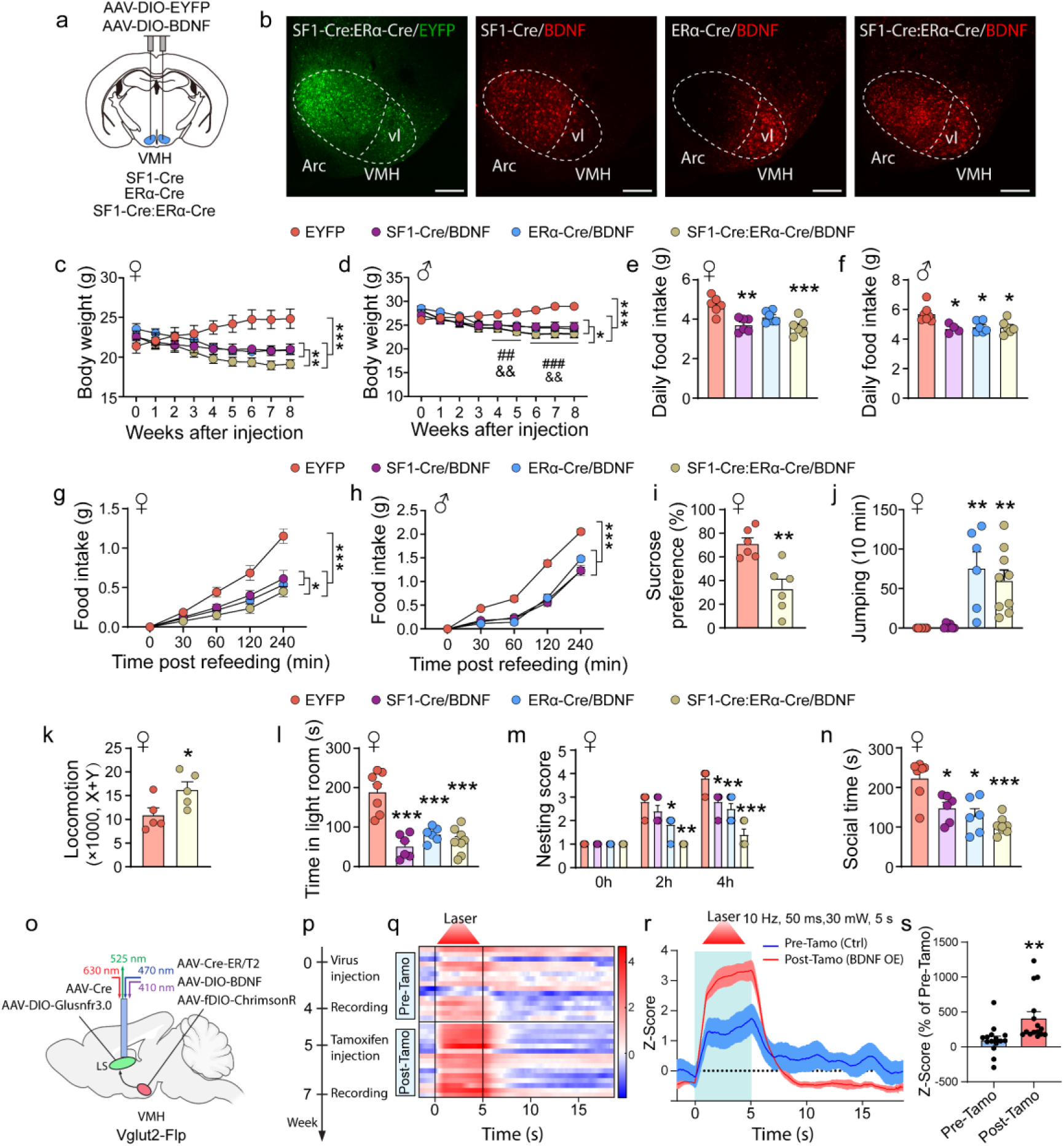
Augmented BDNF action from SF1/ERa neurons leads to anorexia like phenotypes. **(a)** Diagram of viral delivery (AAV-DIO-EYFP, AAV-DIO-BDNF-mCherry) into the VMH of SF1-Cre, ERa-Cre, and SF1-Cre::ERa-Cre mice. **(b)** Representative images showing patterns of EYFP or NaChBac expression in the VMH of SF1-Cre, ERa-Cre, and SF1-Cre::ERa-Cre mice. Scale bar, 250 μm. **(c, d)** Body weight trajectories of female **(c)** and male **(d)** SF1-Cre, ERa-Cre, and SF1-Cre::ERa-Cre mice after VH injection of AAV-DIO-EYFP or AAV-DIO-BDNF. Females, n=6 (EYFP, SF1-Cre+BDNF, ERa-Cre+BDNF), n=12 (SF1-Cre::ERa-Cre+BDNF); males, n=7 (EYFP), n=4 (SF1-Cre+BDNF), n=6 (ERa-Cre+BDNF, SF1-Cre::ERa-Cre+BDNF); two-way ANOVA; *P<0.05, **P<0.01, ***P<0.001. **(e, f)** Daily food intake in EYFP- and BDNF-expressing female **(e)** or male **(f)** mice. females, n=6 per group; males, n=4-7 per group; one-way ANOVA; *P<0.05, **P<0.01, ***P<0.001 versus EYFP. **(g, h)** Cumulative food intake during fasting–refeeding in EYFP- and BDNF-expressing female **(g)** or male **(h)** mice. females, n=6 per group; males, n=4-7 per group; two-way ANOVA; *P<0.05, ***P<0.001. **(i)** Sucrose preference in EYFP- and BDNF-expressing female SF1-Cre::Esr-Cre+BDNF mice. n=6 per group; unpaired two-tailed t test, P=0.002. **(j)** Total number of jumps during the 10-min test in female mice. n=6 (EYFP, SF1-Cre+BDNF, ERa-Cre+BDNF), n=12 (SF1-Cre::ERa-Cre+BDNF); one-way ANOVA; **P<0.01 versus EYFP. **(k)** Quantification of locomotor activity during dark phase (TSE system). n = 5 per group; unpaired two-tailed t test; P=0.042. **(l-n)** Time spent in the light room in light-dark test **(l)**, nesting behavior **(m)**, social time with stranger in three-chamber test **(n)** in female mice. n=6-8 per group; one-way ANOVA **(l, n)**, two-way ANOVA **(m)**; *P<0.05, ***P<0.001 versus EYFP. **(o)** Diagram of in vivo optogenetic stimulation of VMH SF1 and ERα neuronal terminals with simultaneous fiber photometry recording of glutamate sensor signals in downstream lateral septum neurons before and after inducible BDNF overexpression. **(p)** Schematic timeline of fiber photometry recording and tamoxifen injection. **(q)** Heatmap of glutamate release Z score signals of individual trails in response to 5-s optogenetic stimulation. **(r)** Averaged trace of glutamate release Z score signals of lateral septum neurons before, during, and after 5-s stimulation of SF1 and ERa neurons axons in VMH before (blue line) and after inducible BDNF knockout (red line). **(s)** Mean of glutamate release Z score during optogenetic stimulation. n=15 trails per group, unpaired two-tailed t test, P<0.001.

Behavioral analyses further revealed that BDNF overexpression in SF1/ERα neurons reduced sucrose preference (Fig. 6i), indicative of anhedonia. Interestingly, unlike BDNF overexpression in the broader Vglut2 population, which increased locomotor activity in OPT, overexpression in ERα or SF1/ERα neurons did not significantly alter open-field locomotion in either females (Supplementary Figs. S5a and 5b) or males (Supplementary Fig. S5c). However, mice overexpressing BDNF in ERα or SF1/ERα neurons displayed pronounced spontaneous jumping behaviors in both females (Fig. 6j; **Supplementary Video 4**) and males (Supplementary Fig. S5d), which likely confounded locomotor measurements obtained in OFT. Consistent with this possibility, assessment in TSE metabolic home-cage chambers revealed a significant increase in locomotor activity in mice with BDNF overexpression in SF1/ERα neurons (Fig. 6k). In contrast, BDNF overexpression in SF1 neurons alone reduced locomotion (Supplementary Figs. S5a-5c) and failed to induce spontaneous jumping behavior (Fig. 6j; Supplementary Fig. S5d). These observations closely mirror the effects observed following NaChBac-mediated activation of SF1 and ERα neurons (Fig. 2j) and further support a specific role for ERα neurons in mediating the hyperactivity component of the AN-like phenotype.

In addition to alterations in feeding and locomotor activity, female mice overexpressing BDNF in SF1, ERα, or SF1/ERα neurons spent significantly less time in the illuminated compartment in LDT (Fig. 6l) and exhibited impaired nest-building behavior (Fig. 6m), indicative of increased anxiety-like behavior. Similarly, all three groups showed reduced sociability in the three-chamber social interaction assay, with SF1/ERα mice exhibiting the most pronounced deficits (Fig. 6n). Comparable effects were observed in male mice, although BDNF overexpression in ERα neurons alone did not significantly affect anxiety-like behavior or sociability (Supplementary Figs. S5e-5j), suggesting sexually dimorphic contributions of ERα neurons to these behavioral phenotypes.

Previous experiments showed that BDNF deletion markedly reduced glutamate release from SF1/ERα neurons, raising the possibility that enhanced BDNF signaling conversely increases glutamatergic transmission. To test this hypothesis, we delivered a combination of AAV-CreER, AAV-DIO-BDNF-p2A-mCherry, and AAV-fDIO-ChrimsonR to the MBH, together with AAV-Cre-GFP and AAV-DIO-GluSnFR3.0 to the LS of *Vglut2-Flp* mice (Fig. 6o). This strategy enabled tamoxifen-inducible BDNF overexpression in MBH glutamatergic neurons while allowing optogenetic stimulation of their projections and simultaneous monitoring of glutamate release in the LS before and after tamoxifen administration within the same animal (Fig. 6p). Following tamoxifen treatment, optogenetically evoked glutamate sensor responses were markedly enhanced, as illustrated by recordings from individual animals (Fig. 6q), representative traces (Fig. 6r), and quantitative analyses comparing pre- and post-tamoxifen responses (Fig. 6s). These findings demonstrate that increased BDNF signaling potentiates glutamate release from MBH glutamatergic neurons and support a model in which BDNF-driven enhancement of glutamatergic transmission contributes to the emergence of AN-like phenotypes.

Collectively, these findings demonstrate that selective enhancement of BDNF signaling in SF1/ERα neurons and resulting increase in glutamate release are sufficient to reproduce a broad spectrum of AN-like phenotypes, including voluntary restrictive feeding, weight loss, anhedonia, anxiety-like behavior, social avoidance, and hyperactivity. Moreover, the results reveal distinct contributions of SF1 and ERα neuronal populations, with ERα neurons playing a particularly prominent role in mediating hyperactivity and sex-dependent behavioral effects.

### BDNF-induced anorexia phenotypes are rescued by deletion of glutamate release

Our preceding experiments demonstrated that loss of glutamate release completely abolished the AN-like phenotypes induced by chronic activation of SF1/ERα neurons, whereas BDNF deletion produced a delayed but substantial rescue. These findings suggested that BDNF may act upstream of glutamatergic transmission. To test this possibility, we first examined neuronal activity in downstream targets of SF1/ERα neurons following disruption of either glutamate release or BDNF signaling. We assessed c-Fos expression in established downstream targets of MBH SF1/ERα neurons. Chronic activation of SF1/ERα neurons by NaChBac expression markedly increased c-Fos expression in the paraventricular thalamus (PVT), an effect that was completely abolished by CRISPR-mediated deletion of Vglut2 (Figs. 7a and 7b). Similar reductions in c-Fos expression were observed in additional downstream regions, including the lateral septum (LS; Supplementary Figs. S6a and S6b), preoptic area (POA; Supplementary Figs. S6c and S6d), periaqueductal gray (PAG; Supplementary Figs. S6e and S6f), and peri-locus coeruleus region (Peri-LC; Supplementary Figs. S6g and S6h). These findings indicate that glutamatergic transmission from SF1/ERα neurons is required for the activation of multiple downstream targets.

**Figure. 7.**
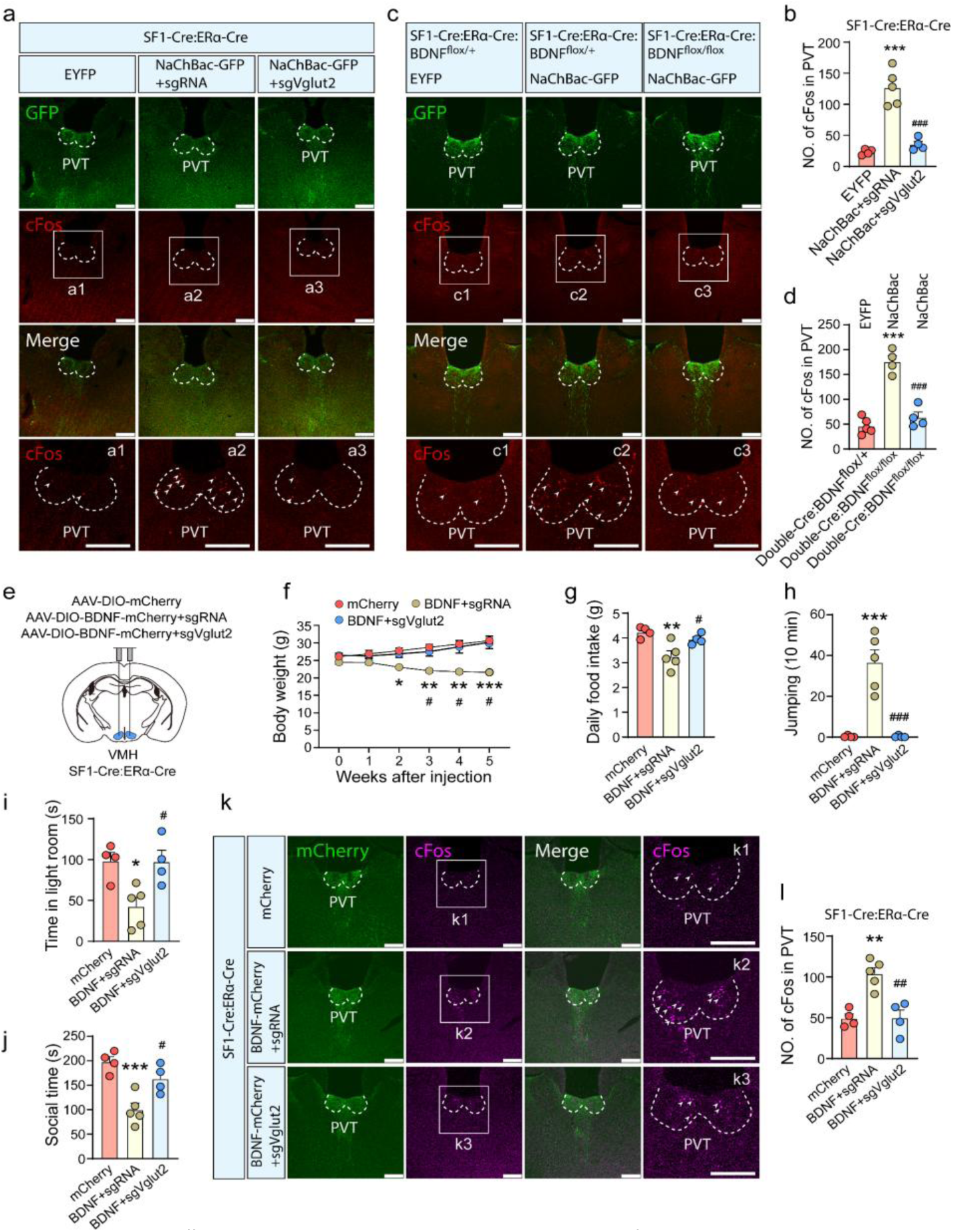
Rescuing effects on BNDF-induced anorexia phenotypes by deletion of glutamate release. **(a)** Representative images showing GFP and c-Fos (arrows) expression patterns in the PVT, a downstream target of the VMH, in AAV-DIO-EYFP, AAV-DIO-NaChBac+sgRNA, AAV-DIO-NaChBac+sgVglut2 injection SF1-Cre::ERa-Cre mice. scale bar=250 um. **(b)** Representative images showing GFP (green) and c-Fos (red, arrows) expression patterns in the PVT, a downstream target of the VMH, in SF1-Cre::ERa-Cre::BDNFflox/++EYFP, SF1-Cre::ERa-Cre::BDNFflox/++NaChBac, SF1-Cre::ERa-Cre::BDNFflox/flox+NaChBac mice. scale bar=250 um. **(c)** Number of c-Fos positive cells in the PVT region related to panel A. n=4-5 per group; one-way ANOVA; ***P<0.001 versus EYFP; ###P<0.001 versus NaChBac+sgRNA. **(d)** Number of c-Fos positive cells in the PVT region related to panel A. n=4-5 per group; one-way ANOVA; ***P<0.001 versus EYFP; ###P<0.001 versus SF1-Cre::ERa-Cre::BDNFflox/flox+NaChBac. **(e)** Diagram of viral delivery (AAV-DIO-mCherry, AAV-DIO-BDNF-mCherry+sgRNA, AAV-DIO-BDNF-mCherry+sgVglut2) into the VMH of SF1-Cre::ERa-Cre mice. **(f)** Body weight trajectories of SF1-Cre::ERa-Cre mice after VH injection of virus. n=4 (EYFP, BDNF+sgVglut2), n=5 (BDNF+sgRNA); two-way ANOVA; *P<0.05, **P<0.01, ***P<0.001 versus mCherry; #P<0.05 versus BDNF+sgRNA. **(g)** Daily food intake. n=4-5 per group, one-way ANOVA, **P=0.006 versus mCherry; #P=0.043 versus BDNF+sgRNA. **(h)** Total number of jumps during a 10-min test. n=4-5 per group; one-way ANOVA; ***P<0.01 versus mCherry; ###P<0.001 versus BDNF+sgRNA. **(i, j)** time spent in the light room in light-dark test (I), social time with stranger in three-chamber test (J). n=4-5 per group; one-way ANOVA; *P<0.05, ***P<0.01 versus mCherry; #P<0.05 versus BDNF+sgRNA. **(k)** Representative images showing mCherry (green) and c-Fos (magenta, arrows) expression patterns in the PVT, a downstream of VMH, in mCherry, BDNF+sgRNA, BDNF+sgVglut2 injection SF1-Cre::ERa-Cre mice. scale bar=250 um. **(l)** number of c-Fos positive cells in PVT region related to panel K. n=4-5 per group; one-way ANOVA; ***P<0.001 versus mCherry; ###P<0.001 versus BDNF+sgRNA.

Interestingly, selective deletion of BDNF produced a similar reduction in c-Fos expression within the PVT (Figs. 7c and S7d), as well as in the LS, POA, PAG, and Peri-LC (Supplementary Fig. S7). Together with our glutamate-release measurements, these data suggest that BDNF romotes activation of downstream circuits by facilitating glutamatergic output from SF1/ERα neurons.

We next asked whether the AN-like phenotypes induced by BDNF overexpression depend on glutamate release. To address this question, we co-injected AAV-DIO-BDNF-p2A-mCherry together with either AAV-DIO-Cas9-sgVglut2 or a control sgRNA vector into the VMH of SF1-Cre::ERα-Cre mice (Fig. 7e). As expected, mice receiving BDNF plus control sgRNA exhibited progressive body weight loss relative to controls (Fig. 7f). Remarkably, deletion of Vglut2 completely prevented this phenotype, resulting in near-normal body weight trajectories (Fig. 7f). Consistent with the restoration of body weight, CRISPR-mediated deletion of Vglut2 abolished the major behavioral effects induced by BDNF overexpression, including voluntary restrictive feeding (Fig. 7g), spontaneous jumping behavior (Fig. 7h; **Supplementary Video 5**), anxiety-like behavior in LDT (Fig. 7i), and reduced sociability (Fig. 7j). These findings indicate that the behavioral and metabolic effects of elevated BDNF signaling require intact glutamatergic transmission.

To further assess circuit-level consequences, we examined c-Fos expression in downstream target regions. Deletion of Vglut2 reversed the BDNF-induced increase in c-Fos expression within the PVT (Figs. 7k and 7l), LS (Supplementary Figs. S8a and S8b), POA (Supplementary Figs. S8c and S8d), PAG (Supplementary Figs. S8e and S8f), and Peri-LC (Supplementary Figs. S8g and S8h). These observations further support the conclusion that enhanced BDNF signaling promotes excitation of downstream circuits primarily through increased glutamate release.

BDNF has been proposed to modulate glutamatergic signaling through both presynaptic enhancement of glutamate release and postsynaptic potentiation of glutamate receptor activity^42^. If BDNF acted predominantly through postsynaptic mechanisms, overexpression of BDNF in SF1/ERα neurons would be expected to increase their intrinsic activity. However, c-Fos expression within SF1/ERα neurons was not elevated by BDNF overexpression compared with mCherry controls or mice receiving both BDNF overexpression and Vglut2 deletion (Supplementary Fig. S9). These findings argue against a major contribution of postsynaptic glutamate receptor potentiation and instead support a presynaptic mechanism whereby BDNF enhances glutamate release.

Given the critical role of the MBH in feeding regulation, we next sought to identify downstream pathways that might mediate the observed AN-like phenotypes. Previous studies implicated MBH projections to trigeminal mesencephalic nucleus (Me5) neurons in the control of jaw movement and food ingestion^43^. We therefore examined activity within the peri-LC region, which contains Me5 neurons. Mapping studies revealed that the strongest c-Fos induction occurred in the laterodorsal tegmental nucleus (LDTg), whereas only minimal c-Fos expression was detected in Me5 neurons in either NaChBac-expressing (Supplementary Figs. S6g and S6h) or BDNF-overexpressing mice (Supplementary Figs. S7g and S7h). To directly test the contribution of Peri-LC neurons, we combined activation of SF1/ERα neurons with transsynaptic tracing and targeted silencing of downstream Peri-LC neurons. Specifically, AAV-DIO-NaChBac-GFP and AAV-DIO-WGA-GFP were co-injected into the VMH of *SF1-Cre::ERα-Cre* mice, followed by delivery of AAV-Flp-DOG to the Peri-LC. This strategy enabled selective labeling of synaptically connected Peri-LC neurons. Concurrent delivery of AAV-fDIO-Cre together with either AAV-DIO-Kir2.1-dTomato or control AAV-DIO-mCherry allowed selective silencing of these traced neurons (Supplementary Fig. S10a).

We confirmed robust expression of Kir2.1 within traced Peri-LC neurons and found that Kir2.1 effectively suppressed the elevated c-Fos expression induced by NaChBac-mediated activation of SF1/ERα neurons (Supplementary Figs. S10b and S10c), indicating successful neuronal inhibition. Despite this effective silencing, inhibition of Peri-LC neurons failed to rescue either the body weight loss (Supplementary Fig. S10d) or lethality (Supplementary Fig. S10e) associated with chronic activation of SF1/ERα neurons.

Collectively, these findings demonstrate that BDNF-induced AN-like phenotypes are mediated through enhanced glutamatergic transmission and downstream circuit activation. Furthermore, while Peri-LC neurons are activated by SF1/ERα neurons, they are not a major contributor to the pathological feeding and body weight phenotypes observed in this model. These results support a model in which BDNF acts upstream of glutamate release to potentiate excitatory output from SF1/ERα neurons, thereby driving the behavioral and metabolic manifestations of AN-like pathology.

### Refractory anorexic effects by activation of MBH glutamatergic neurons

AN remains one of the most difficult psychiatric disorders to treat, highlighting the need for a deeper mechanistic understanding of the neural circuits underlying pathological food restriction^44^. To determine whether the anorectic phenotype induced by chronic activation of MBH glutamatergic neurons is reversible, we tested whether it could be overcome by genetic manipulations known to drive robust positive energy balance and hyperphagia. In previous studies, chronic NaChBac-mediated activation of AgRP neurons produced profound hyperphagia and severe obesity^37^. We therefore asked whether activation of this potent orexigenic neuronal population could counteract the voluntary starvation induced by activation of MBH glutamatergic neurons. To this end, we generated *AgRP-Cre::Vglut2-Cre* mice and delivered AAV-DIO-NaChBac-GFP bilaterally to both the arcuate nucleus and VMH (Fig. 8a). Correct viral targeting and robust neuronal activation were verified by c-Fos expression in both AgRP and MBH glutamatergic neurons (Fig. 8b).

**Fig. 8.**
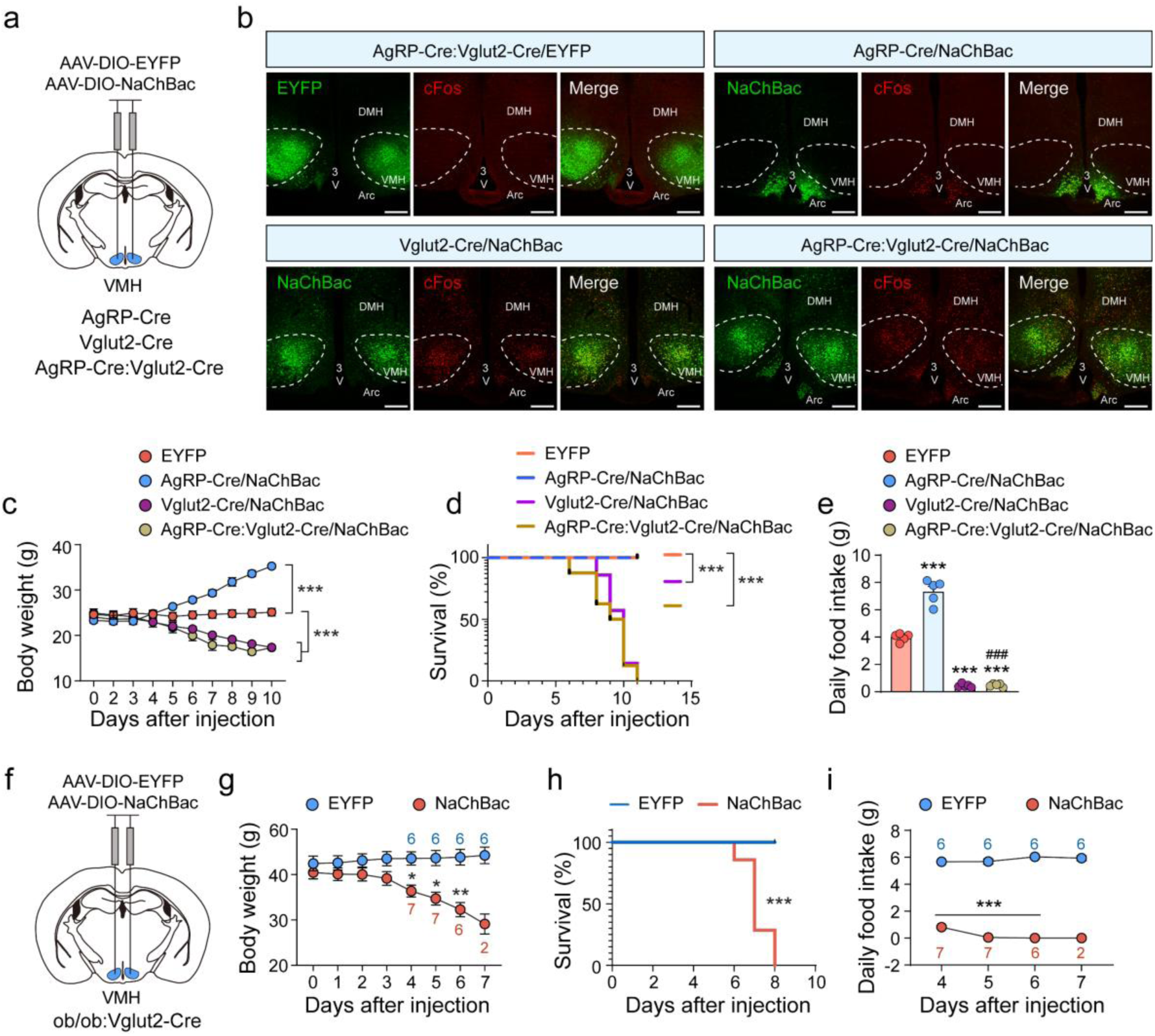
Refractory anorexic effects by activation of ventral hypothalamic glutamatergic neurons. **(a)** Diagram of viral delivery (AAV-DIO-EYFP, AAV-DIO-NaChBac) into VH of AgRP-Cre, Vglut2-Cre, and AgRP-Cre::Vglut2-Cre. **(b)** Representative images showing EYFP or NaChBac expression patterns in the VMH, and robust c-Fos induction (arrows) in NaChBac-expressing mice. Scale bar, 250 μm. **(c)** Body weight trajectories of AgRP-Cre, Vglut2-Cre, AgRP-Cre::Vglut2-Cre mice after EYFP or NaChBac injection. n=8 (EYFP), n=6 (AgRP-Cre), n=8 (Vglut2-Cre), n=7 (AgRP-Cre::Vglut2-Cre); two-way ANOVA; ***P<0.001. **(d)** Kaplan–Meier survival analysis of AgRP-Cre, Vglut2-Cre, AgRP-Cre::Vglut2-Cre mice after EYFP or NaChBac injection. Mice were considered as non-survival when body weight loss reached 30%, log-rank test, ***P<0.001 versus EYFP. **(e)** Daily food intake. n=5 per group; one-way ANOVA, ***P<0.001 versus EYFP, ^###^P<0.001 versus AgRP-Cre/NaChBac. **(f)** Diagram of viral delivery (AAV-DIO-EYFP, AAV-DIO-NaChBac) into VH of ob/ob::Vglut2-Cre mice. **(g)** Body weight trajectories of ob/ob::Vglut2-Cre mice after EYFP or NaChBac injection. n=6 (EYFP), n=7 (NaChBac); two-way ANOVA; *P<0.05, **P<0.01. **(h)** Kaplan–Meier survival analysis of ob/ob::Vglut2-Cre mice after EYFP or NaChBac injection. Mice were considered non-survival when body weight loss reached 30%, log-rank test, ***P<0.001. **(i)** Daily food intake. n=6-7 per group; two-way ANOVA, ***P<0.001.

As expected, activation of AgRP neurons alone produced rapid body weight gain, whereas activation of Vglut2-expressing neurons alone resulted in marked weight loss (Fig. 8c). Strikingly, simultaneous activation of both AgRP and Vglut2 neurons failed to rescue the anorectic phenotype. Instead, these mice closely phenocopied animals in which only Vglut2 neurons were activated, exhibiting progressive body weight loss and eventual lethality (Figs. 8c and 8d). Furthermore, mice with concurrent activation of AgRP and Vglut2 neurons displayed a degree of voluntary food restriction indistinguishable from that observed following activation of Vglut2 neurons alone (Fig. 8e). These findings indicate that the powerful orexigenic drive generated by AgRP neurons is insufficient to overcome the voluntary starvation induced by chronic activation of MBH glutamatergic neurons.

To further assess the robustness of this anorectic state, we examined whether genetically driven hyperphagia resulting from leptin deficiency could reverse the phenotype. For this purpose, *Vglut2-Cre* mice were crossed with leptin-deficient *ob/ob* mice, a model characterized by profound hyperphagia and severe obesity. AAV-DIO-NaChBac-GFP or control AAV-DIO-GFP vectors were then delivered to the MBH of *Vglut2-Cre::ob/ob* mice (Fig. 8f). As expected, control GFP-injected mice exhibited progressive weight gain and developed severe obesity (Fig. 8g). In contrast, activation of MBH glutamatergic neurons in *ob/ob* mice induced a rapid and progressive loss of body weight despite the absence of leptin signaling (Fig. 8g). Remarkably, all NaChBac-expressing mice succumbed within eight days of viral delivery (Fig. 8h), recapitulating the lethal phenotype observed in *Vglut2-Cre* mice with intact leptin signaling. Consistent with the marked weight loss, these mice exhibited a profound reduction in food intake relative to controls (Fig. 8i), indicating that voluntary food restriction remained the primary driver of the phenotype.

Collectively, these findings demonstrate that the anorectic state induced by chronic activation of MBH glutamatergic neurons is highly refractory to physiological mechanisms that normally promote feeding. Neither AgRP neuron activation nor leptin deficiency, two of the most potent drivers of hyperphagia and positive energy balance, was sufficient to overcome the voluntary starvation induced by this manipulation. These results suggest that activation of MBH glutamatergic neurons engages a dominant neural program that suppresses feeding behavior even in the presence of strong homeostatic signals promoting food intake.

## Discussion

Progress in understanding the neural basis of anorexia nervosa (AN) has been hindered by the lack of animal models that faithfully recapitulate the core clinical features of the disorder^12,45^. The most widely used model, activity-based anorexia (ABA), relies on imposed food restriction and involuntary starvation to generate a negative energy state, a paradigm fundamentally distinct from the voluntary food restriction that drives AN pathogenesis in humans^12,46^. Consequently, existing models capture only selected aspects of the disorder and have provided limited insight into the neural mechanisms underlying AN. Similarly, BDNF-based genetic models, although motivated by human genetic findings, rely on fasting paradigms and exhibit variable AN-associated phenotypes ^21^. Other models, including the *anx/anx* mouse, display self-starvation but fail to reproduce additional hallmark symptoms such as hyperactivity ^47^. In contrast, chronic activation of MBH SF1/ERα neurons or selective enhancement of BDNF signaling within these neurons produced voluntary food restriction, hyperactivity, anxiety-like behavior, anhedonia, social avoidance, and enhanced sensitivity to environmental stressors, collectively capturing the core behavioral manifestations of AN.

Beyond reproducing core AN-like phenotypes, our model possesses several features that increase its translational relevance. First, it directly implicates BDNF, a molecule strongly linked to AN through human genetic studies. Second, the involvement of ERα-expressing neurons provides a potential mechanistic explanation for the markedly higher prevalence of AN in females. Third, the exaggerated jumping behavior elicited by exposure to novel environments resembles the well-established association between AN onset and stressful or traumatic life experiences in humans. Together, these findings establish a novel experimental model that recapitulates the central features of AN while identifying VMH SF1/ERα neurons as a key neural substrate capable of driving AN-like pathophysiology.

The VMH has historically been regarded as a “satiety center” because lesions of this region produce hyperphagia and obesity^48,49 50^. Consistent with this view, acute activation and inhibition of VMH SF1 neurons suppress and promote feeding, respectively^51^. Beyond feeding regulation, VMH neurons have also been implicated in defensive responses, fear-related behaviors, and social interactions^31,52–54^. Within the VMH, ERα-expressing neurons partially overlap with SF1 neurons and contribute to sexually dimorphic regulation of feeding, anxiety-related behavior, and locomotion^29,55,56,30^. Collectively, these observations position VMH neurons as important regulators of behavioral domains that are profoundly disrupted in AN. However, previous studies have largely relied on acute manipulations or loss-of-function approaches and therefore provide limited insight into the consequences of chronic neuronal dysregulation, a feature likely to underlie many neuropsychiatric disorders, including AN^33^. By chronically activating SF1/ERα neurons using NaChBac-mediated depolarization, our study reveals that persistent elevation of VMH neuronal activity is sufficient to induce a syndrome strikingly reminiscent of AN, thereby extending previous work from normal physiological regulation to disease-relevant mechanisms.

Our findings further identify distinct contributions of SF1 and ERα neuronal populations to individual components of the AN phenotype. Although activation of either population contributed to voluntary food restriction, anxiety-like behavior, and reduced sociability, the effects on locomotion diverged markedly. Chronic activation of SF1 neurons reduced locomotor activity, whereas activation of ERα neurons increased locomotion. The locomotor suppression associated with SF1 neuron activation is consistent with previous evidence linking these neurons to the regulation of skeletal muscle metabolism^55^. In contrast, ERα neurons have been shown to promote physical activity, particularly in females^29^. Our data suggest that chronic activation of ERα neurons exerts a dominant pro-locomotor effect that overrides the locomotor suppression produced by SF1 neurons, resulting in the hyperactivity observed following combined SF1/ERα activation. These findings identify ERα neurons as a critical component of the circuitry underlying AN-associated hyperactivity. Notably, activation of ERα neurons produced stronger anxiety-like and social deficits in females than in males. Similarly, BDNF overexpression within ERα neurons elicited robust behavioral phenotypes in females but comparatively modest effects in males. These observations reveal a sex-dependent contribution of ERα neurons to AN-like behaviors and may provide a mechanistic framework for understanding the markedly greater incidence of AN in adolescent females. Given the profound hormonal changes associated with puberty and the established role of estrogen signaling in VMH function^57^, altered activity of ERα neurons may contribute to the heightened vulnerability of females to AN.

A major finding of the current study is the identification of BDNF as a critical mediator of SF1/ERα neuron-driven AN-like phenotypes. Deletion of BDNF markedly attenuated the behavioral and metabolic consequences of chronic SF1/ERα activation, whereas BDNF overexpression recapitulated many of the same phenotypes. These findings are consistent with previous studies demonstrating that disruption of hypothalamic BDNF signaling promotes obesity, a phenotype opposite to that observed here^43,58–60 58,61^. However, reports of circulating and central BDNF levels in AN patients have yielded conflicting results, with studies reporting increases, decreases, or no significant changes^62–64^. The present findings suggest that such inconsistencies may reflect heterogeneity in the underlying mechanisms responsible for disease development. Defects occurring upstream of BDNF signaling may lead to compensatory increases in BDNF expression, whereas abnormalities downstream of BDNF action may produce reductions through feedback regulation. Interestingly, the AN-associated BDNF^Val66Met^ variant is generally considered to impair BDNF function^65^, a finding that appears difficult to reconcile with our observation that enhanced BDNF signaling promotes AN-like phenotypes. Resolving this apparent discrepancy will require direct examination of the consequences of BDNF^Val66Met^ expression specifically within SF1/ERα neurons.

Mechanistically, our data support a model in which BDNF acts upstream of glutamatergic transmission to drive AN-like behaviors. Selective disruption of glutamate release completely rescued the behavioral and metabolic abnormalities induced by chronic SF1/ERα activation, indicating that glutamate serves as the principal downstream effector. Consistent with this conclusion, chronic activation of SF1/ERα neurons increased c-Fos expression throughout multiple downstream target regions, and these effects were abolished by deletion of Vglut2. Loss of BDNF produced a similar reduction in downstream neuronal activation and substantially reduced glutamate release, whereas BDNF overexpression enhanced glutamatergic signaling and increased activity throughout downstream circuits. Although BDNF has been reported to regulate glutamate transmission^42^, its role in maintaining basal glutamate release from VMH neurons has not been previously demonstrated. Furthermore, BDNF overexpression failed to increase c-Fos expression within SF1/ERα neurons themselves, arguing against a dominant postsynaptic mechanism. Instead, our findings support a model in which BDNF potentiates presynaptic glutamate release, thereby amplifying excitatory output from SF1/ERα neurons and driving AN-like behavioral states.

An especially striking observation was the remarkable resistance of the starvation phenotype to powerful orexigenic manipulations. Whereas deletion of either BDNF or glutamate release rescued the phenotype, neither chronic activation of AgRP neurons nor leptin deficiency, two manipulations known to induce profound hyperphagia and obesity, was able to reverse the voluntary starvation elicited by activation of MBH glutamatergic neurons. These findings suggest that the induced anorectic state overrides endogenous homeostatic feeding circuits even under conditions of extreme hunger. Such dominance may help explain the clinical difficulty of treating AN through interventions aimed solely at increasing caloric intake ^66^. We also examined the potential contribution of VMH projections to the trigeminal mesencephalic nucleus (Me5), which have previously been implicated in the regulation of jaw movements and food intake^43^. However, we observed minimal activation of Me5 neurons following either SF1/ERα activation or BDNF overexpression, and chronic inhibition of Peri-LC neurons failed to rescue the starvation phenotype. These findings argue against a major contribution of the Me5 pathway in mediating the restrictive feeding observed in our model. Instead, SF1/ERα neurons project broadly to regions including the PVT, PAG, BNST, amygdala, and striatum^67–69 31,67,68,70,71^, many of which regulate feeding, anxiety, motivation, and social behavior. Importantly, increased activity within several of these regions has been reported in AN patients^72–74^, suggesting that the circuits identified here may be relevant to human disease.

From an evolutionary perspective, it is remarkable that chronic activation of such a small neuronal population can drive self-imposed starvation to the point of lethality, despite the existence of highly redundant feeding circuits designed to preserve survival^75^. One possibility is that these neurons engage a self-reinforcing circuit that progressively strengthens restrictive feeding behavior. In this regard, physical activity and exercise are known to increase BDNF expression in the VMH^76^. Our data demonstrate that enhanced BDNF signaling promotes both hyperactivity and food restriction, raising the intriguing possibility of a positive feedback loop in which increased activity elevates BDNF signaling, which in turn further promotes hyperactivity and restrictive feeding. Future studies will be needed to directly test this hypothesis.

In summary, we establish a novel animal model that faithfully recapitulates the central behavioral features of AN, including voluntary self-starvation, hyperactivity, anxiety-like behavior, social avoidance, and female-biased susceptibility. Our findings identify VMH SF1/ERα neurons as a critical neural substrate for AN-like pathogenesis and reveal enhanced BDNF-dependent glutamatergic transmission as a key mechanistic driver of these phenotypes. These results provide new insight into the neural circuitry underlying AN and identify potential targets for therapeutic intervention.

## Acknowledgements

The authors thank members in the Tong lab for inputs and discussion. Q.T. is supported by DOD HT94252310156 and R01DK109934 (QT and BRA), NIH R01 DK136284, R01 DK 135212 and R01 DK 131466 (QT), and R01MH139750 (YX). The project is benefited from the Optogenetics and Viral Vectors Core, supported by NIH IDDRC grant 1 U54 HD083092, and the Gene Vector Core, at Baylor College of Medicine for providing viral preparations. QT is the holder of the Cullen Chair in Molecular Medicine and Hans J. Eberhard MD, PhD and Irma Gigli, MD Distinguished Chair in Immunology at McGovern Medical School.

## Author contribution

The experiments were mainly conducted by C.S. with helps from Y.X., M.Y., D.L., Y.C., and R.Y. B.R.A. provided essential reagents. C.S. wrote the manuscript with significant inputs from B.R.A., and Q. T.

## EXPERIMENTAL MODEL AND STUDY PARTICIPANT DETAILS

### Animals

Animal care and experimental procedures were approved by the Institutional Animal Care and Use Committee of the University of Texas Health Science Center at Houston. Mice were maintained at 21–22 °C under a 12-h light/12-h dark cycle with standard pellet chow (LabDiet 5053; 20% protein diet and 4.5%; 3.02 kcal/g) and water provided ad libitum, unless otherwise specified for fasting experiments. Animals were group-housed under standard conditions, and singly housed when measuring daily food intake or when placed in metabolic cages. Vglut2-Cre, Vglut2-Flp, SF1-Cre, AgRP-Cre, ob/ob, Esr1-Cre (ERα-Cre) mice were purchased from The Jackson Laboratory. BDNF^flox/flox^ mice were kindly provided by Dr. Baoji Xu from U. Florida. For all experiments, mice from the same litters were randomly assigned to different treatment groups and were 6–8 weeks of age at the time of surgical procedures.

### Stereotaxic Surgery

Stereotaxic surgeries for viral delivery were performed as previously described. Briefly, mice were anesthetized with a ketamine/xylazine cocktail (100 and 10 mg/kg, respectively), and their heads were secured in a stereotaxic frame. Viral vectors were delivered through a 0.5 mL syringe (Neuros Model 7000.5 KH, point style 3; Hamilton, Reno, NV, USA) mounted on a motorized stereotaxic injector (Quintessential Stereotaxic Injector; Stoelting, Wood Dale, IL, USA) at a rate of 30 nL/min. Viral preparations were titered to ∼10¹² particles/ml. The stereotaxic coordinates were as follows: VMH, anteroposterior (AP) −1.50 mm, mediolateral (ML) ±0.35 mm and ±0.70 mm, dorsoventral (DV) −5.60 mm (60 nL/site); LS, AP +0.50 mm, ML ±0.50 mm, DV −3.70 mm (DV −3.50 mm for fiber implantation, 100 nL/site); Me5, AP −5.35 mm, ML ±0.90 mm, DV −3.45 mm (100 nL/site).

For NaChBac or BDNF overexpression, AAV-EF1α-DIO-EYFP(Addgene_27056), AAV-EF1α-DIO-NaChBac-GFP (Baylor Gene Vector Core), AAV-EF1α-DIO-mCherry (Addgene_50459), or AAV-EF1α-DIO-BDNF-mCherry (Baylor Gene Vector Core) was bilaterally injected into the VMH.

For Vglut2 knockdown, AAV-EF1α-DIO-NaChBac-GFP or AAV-EF1α-DIO-BDNF-mCherry was mixed (1:1) with either AAVDJ8-DIO-SaCas9-U6-sgRNA or AAVDJ8-DIO-SaCas9-U6-sgVglut2 (Baylor Gene Vector Core) and bilaterally injected into the VMH.

To measure glutamate release in LS neurons following activation of upstream VMH neurons in the Vglut2 knockdown model, AAVDJ8-DIO-SaCas9-U6-sgRNA or AAVDJ8-DIO-SaCas9-U6-sgVglut2 was co-injected with AAV-Syn-DIO-ChrimsonR-tdTomato (Addgene_62723, 1:1) into the bilateral VMH; AAV-Cre-GFP and AAVDJ8-hSyn-GluSnFR3.0 (Baylor Gene Vector Core, 1:1) were co-injected into the bilateral LS of SF1-Cre:ERα-Cre mice. To measure glutamate release in LS neurons following activation of upstream VMH neurons in the inducible BDNF knockout model, AAVDJ8-hSyn-Cre-ER/T2 (Addgene_178059), AAV-EF1α-DIO-BDNF-mCherry, AAV-EF1α-Flp (Baylor Gene Vector Core), and AAV-EF1α-fDIO-ChrimsonR-tdTomato (Addgene_128589) were mixed at a 1:1:1:1 ratio and bilaterally injected into the VMH; AAV-Cre and AAVDJ8-hSyn-GluSnFR3.0 (1:1) were co-injected into the bilateral LS of BDNF^flox/flox^ mice. To measure glutamate release in LS neurons following activation of upstream VMH neurons in the inducible BDNF overexpression model, AAVDJ8-hSyn-Cre-ER/T2, AAV-EF1α-DIO-BDNF-mCherry, and AAV-EF1α-fDIO-ChrimsonR-tdTomato were mixed at a 1:1:1 ratio and bilaterally injected into the VMH; AAV-Cre and AAVDJ8-hSyn-GluSnFR3.0 (1:1) were co-injected into the bilateral LS of Vglut2-Flp mice. After viral injection, we implanted the fiber-optic cannular (Ø1.25 mm, Ø400 μm Core, 0.39 NA; Doric Lens) 0.20 mm higher above the LS.

For inhibition of VMH downstream neurons in the periLC, AAVDJ8-CAG-DIO-WGA-GFP (Canadian Neurophotonics Platform, SCR_016477) and AAV-EF1α-DIO-NaChBac-GFP were mixed (1:1) and bilaterally injected into the VMH. AAV1-EF1α-Flp-DOG-NW (Addgene_75469), AAV9-EF1α-fDIO-Cre (Addgene_121675), and pAAV-EF1α-DIO-Myc-Kir2.1-tdTomato (Baylor Gene Following surgery, mice received carprofen (Rimadyl; Zoetis Inc., Kalamazoo, MI, USA; 5 mg/kg, i.p.) once daily for three consecutive days for postoperative analgesia.

### Inducible BDNF knockout and overexpression

To induce CreERT2-mediated deletion or overexpression of BDNF, tamoxifen (Thermo Scientific, J63509.03) was first dissolved in 10% DMSO/corn oil at a concentration of 50 mg/ml and then diluted with corn oil to a final concentration of 5 mg/ml for injection. Mice received intraperitoneal injections of tamoxifen (50 mg/kg) once daily for five consecutive days.

### Fiber Photometry

In vivo glutamate recordings were performed using the R821/FR-21 Tricolor Multichannel Fiber Photometry system (RWD life science, Shenzhen, China) in an open-top home cage. All tests were conducted during the light cycle following a minimum 4 weeks post-surgery. Mice were habituated in the recording cage for 10 minutes with the optic patch fiber connected, and the mice ID was blinded when the recordings were conducted. Briefly, 470- and 410-nm laser beams were first launched into the fluorescence cube and then coupled into the optical fibers; the 410-nm laser served as amotion-control reference. Glusnfr signal and control emission fluorescence was collected by the camera at the sample rate of 20 Hz. After habituation, Glusnfr signal was recorded. At 1 min after the start of recording, photostimulation was delivered for 5 s using a 630 nm laser source of red light (RWD life science) at a frequency of 10 Hz and a power intensity of 30 mW.

The photometry signal (F) was calculated as the Z score, which is analyzed automatically through the RWD software.

### Body Weight Measurement

Body weight was measured daily in mice injected with AAV-DIO-NachBac and weekly in mice injected with AAV-DIO-BDNF following surgery under a standard chow diet.

### Food Intake Measurement

Daily food intake was measured beginning 3 days after surgery in NachBac-injected Vglut2-Cre mice, 1 week after surgery in NachBac-injected SF1-Cre, ERa-Cre, SF1-Cre::ERa-Cre mice, and 3 weeks after surgery in the BDNF viral injected mice. Mice were singly housed 3 days before the start of food intake measurements. Food intake was recorded for at least 3 consecutive days, and the average value was used as the daily food intake.

### Fasting Refeeding Assay

Fasting refeeding experiments were performed after the completion of food intake measurements. Mice were fasted from 9:00am to 18:00pm, after which standard chow pellets were provided. Food intake was recorded at the 30-, 60-, 120-, and 240-min time points post refeeding.

### Open Field Test (OFT)

Mice were placed individually into an open-field chamber, and their movements were recorded for 15 min using overhead cameras. Locomotion parameters were quantified with EthoVision XT software (Noldus Information Technology).

### Light-Dark box Test (LDT)

The anxiety-like behavior was assessed using the light–dark box test. Mice were initially placed in the dark compartment, and their movement within the light and dark chambers was recorded using an overhead camera and analyzed with the EthoVision XT software (Noldus Information Technology). The time spent in the light compartment was used as an index of anxiety-like behavior, with increased light-chamber exploration reflecting reduced anxiety.

### Nest-building Behavior

Nest-building behavior was assessed after mice had been housed individually for one week. Approximately 2.5 g of a compressed white cotton nesting material piece was placed in each cage at ∼09:00, and nests were evaluated in 4 h. The nest quality was scored on a 5-point scale: 1, no nest attempted; 2, poor nest, with incomplete use of nesting material and minimal structure; 3, fair nest, with full use of nesting material but lacking organized walls; 4, good nest, with full use of nesting material and low but discernible walls; 5, excellent nest, with full use of nesting material and well-defined high walls.

### Three-chamber Sociability Test

The three-chamber test is used to evaluate sociability. The three-chamber apparatus (60 cm × 42.5 cm × 22 cm) was constructed of clear polycarbonate wall and grey Komatex floor, with three equally divided chambers (42.5 cm × 20 cm × 22 cm). Chambers were connected by sliding doors (5.5 cm × 18 cm) in two dividing walls, which control access to left and right chambers. Two stainless-steel barred cups were put in the left and right chambers, besides the walls. During the initial habituation phase, the mice were placed in the middle chamber, and freely explore each of the three chambers with empty stainless-steel barred cups. Immediately after this, an unfamiliar same sex same age mouse which had no previous interaction with the subject mouse, was enclosed in a wire cup. The opposite chamber contained a similar empty wire cup, allowing the subject mouse to explore for 10 min freely. The interaction duration with the stranger mice and empty wire cage was recorded.

### Sucrose Preference Test (SPT)

The sucrose preference test was used to assess anhedonia. Mice were singly housed for one week with ad libitum access to food and water prior to the start of the test. Over the subsequent 3-day adaptation period, each mouse was given two bottles: one containing tap water and the other containing 1% sucrose solution. On the day before testing, mice were fasted from water with food provided *ad libitum*. At 8:00 a.m. on the test day, both bottles were provided, and their positions were switched at 1:00 p.m. Bottle weights were recorded at 8:00 a.m. and 6:00 p.m. Sucrose preference was calculated as: Sucrose preference (%) = [sucrose intake / (sucrose intake + water intake)] × 100%.

### Metabolic Cages Test

Mice were individually housed in chambers of the PhenoMaster cages (TSE systems, Chesterfield, Missouri, USA) with *ad libitum* access to a normal chow diet and water. Locomotion activity levels were measured using indirect calorimetry continuously at different time points for a duration of 5-7 days. Data was averaged through different time points in light and dark cycles, respectively, for comparison. The data from the first day and the last day was removed.

### Perfusion and Tissue Dissection

Mice were anaesthetized with ketamine/xylazine (100 and 10 mg kg⁻¹, respectively). After loss of the pedal reflex is confirmed, animals were transcardially perfused with 15 ml of 0.9% saline followed by 15 ml of 10% formalin. Brains were collected, post-fixed overnight in 10% formalin at room temperature and cryoprotected in 30% sucrose in PBS. Coronal sections (30 µm) were cut on a freezing microtome and stored in PBS containing 0.1% sodium azide at 4 °C.

### Immunostaining and Imaging

Brain sections were washed three times for 5 min each in PBS containing 0.1% Triton X-100, followed by blocking in PBS with 0.1% Triton X-100 and 5% normal donkey serum for 1 h at room temperature. Sections were then incubated overnight at 4 °C in primary antibody solution (primary antibody, 5% normal donkey serum, 0.1% Triton X-100 in PBS). The following primary antibodies were used: rabbit anti-cFos (1:1000, #2250, Cell Signaling Technology), rabbit anti-BDNF(1:200, ANT-010, Alomone Labs), goat anti-GFP (1:1000, 51370, ROCKLAND), rabbit anti-ER alpha (1:500, NBP1-84827), chicken anti-mCherry (1:500, MCHERRY-0020, aves labs).The next day, sections were rinsed three times in PBS with 0.1% Triton X-100 and incubated with secondary antibodies prepared in the same diluent (secondary antibody, 5% normal donkey serum, 0.1% Triton X-100 in PBS). Floating sections were mounted onto glass slides and coverslipped with Fluoromount (Diagnostic BioSystems Inc., Sigma-Aldrich). Images were acquired using a Leica TCS SP5 confocal microscope (Leica Microsystems, Wetzlar, Germany). The fluorescence-positive neurons were recognized and quantified in the software QuPath 0.4.2 (https://qupath.github.io).

### Quantification and Statistical Analysis

GraphPad Prism 9.5.1 (GraphPad Software, Inc., La Jolla, CA, USA) was used for all statistical analyses and construction parts of the figures. Comparisons were made by unpaired two-tailed Student’s t-test, or one-way or two-way ANOVA followed by Tukey’s multiple-comparison post hoc test. Error bars in graphs represent ± s.e.m. All statistical analyses and sample sizes are indicated in the corresponding figure legends.

---

**Figure S1.**
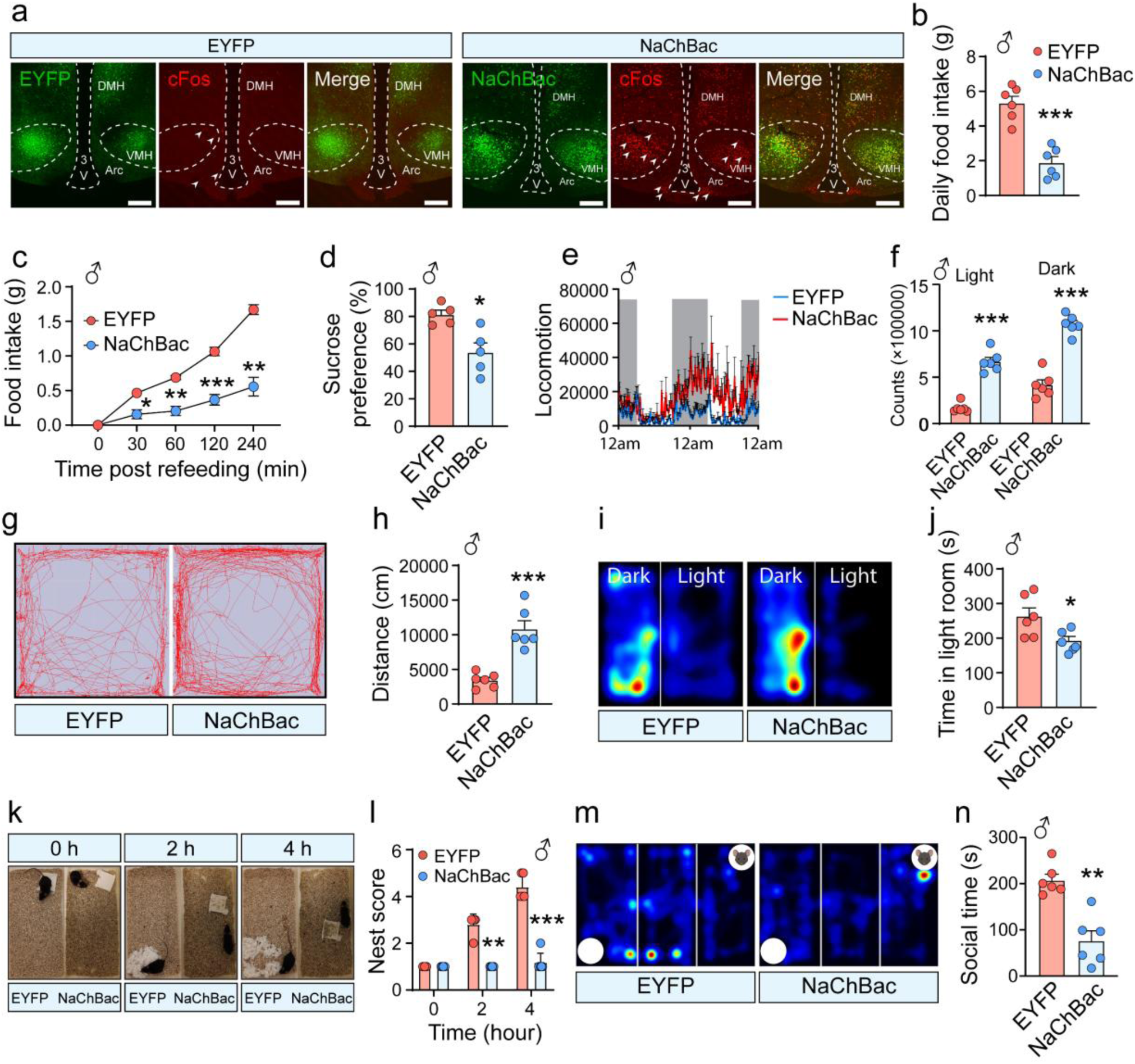
Chronic activation of VH glutamatergic neurons induces typical anorexia-like phenotypes in males (related to Fig.1). **(a)** Representative images showing EYFP or NaChBac expression pattern in VH of Vglut2-Cre mice, and robust c-Fos induction (arrows) in NaChBac-expressing mice. Scale bar=300 μm. **(b)** Daily food intake in EYFP- and NaChBac-expressing male Vglut2-Cre mice. n = 6 per group; unpaired t test; ***P < 0.001. **(c)** Cumulative food intake during fasting–refeeding in EYFP- and NaChBac- expressing male Vglut2-Cre mice. n = 7 (EYFP), n=5 (NaChBac); two-way ANOVA; *P<0.05, **P<0.01, ***P<0.001. **(d)** Sucrose preference in EYFP- and NaChBac-expressing male mice. n = 5 per group; unpaired two-tailed t test, P=0.007. **(e)** Real-time of spontaneous locomotor activity of male mice measured in metabolic cages (TSE). **(f)** Quantification of locomotor activity during light and dark phases. n = 6 per group; unpaired two-tailed t test; ***P<0.001. **(g, h)** Open-field test. **(g)** Representative movement trajectories. **(h)** Total distance traveled in 15 min. n = 6 per group; unpaired two-tailed t test, P<0.001. **(i, j)** Light-dark box test. **(i)** Representative movement trajectories. **(j)** Time spent in the light room. n =6 per group; unpaired two-tailed t test, P=0.028. **(k, l)** Nesting behavior. **(k)** Representative images. **(l)** Nesting scores over time. n = 6 per group; two-way ANOVA; **(m, n)** Three-chamber sociability test. **(m)** Representative trajectories. **(n)** Time spent interacting with a stranger mouse. n = 5 per group; unpaired two-tailed t test; P=0.001.

**Figure S2.**
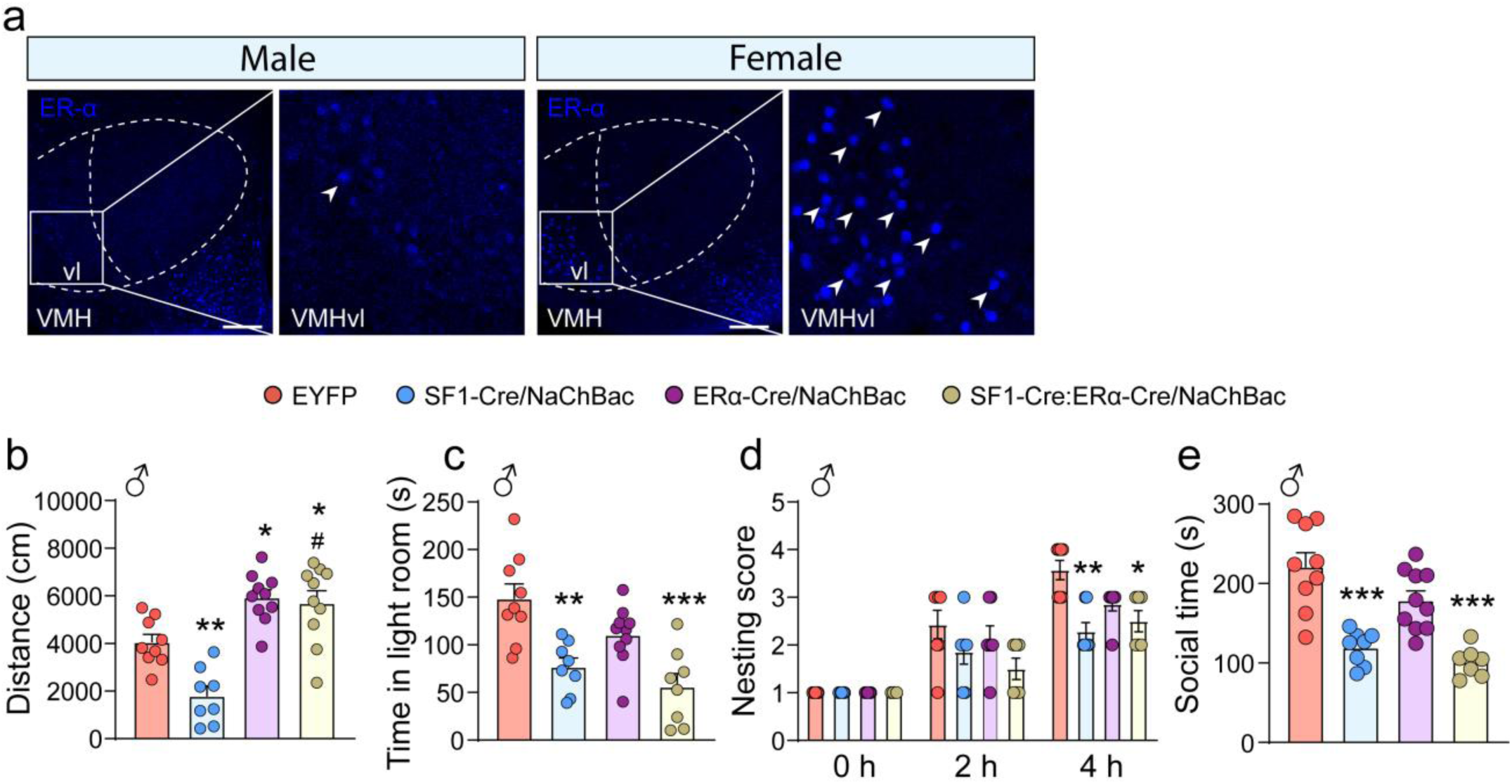
Chronic activation of VH SF1 and ERa neurons in male mice led to typical anorexia phenotypes (related to Fig.2). **(a)** Representative immunostaining images showing ERα expression (arrows) pattern in VMH region in males and females. VMH: ventromedial hypothalamus; VMHvl: ventrolateral subregion of the ventromedial hypothalamus. Scale bar= 250 μm. **(b)** Total distance traveled in 15 min in SF1-Cre::ERa-Cre/EYFP, SF1-Cre/NaChBac, ERa-Cre/NaChBac, SF1-Cre::ERa-Cre/NaChBac male mice. n=8-10 per group; one-way ANOVA; *P<0.05, **P<0.01 versus EYFP; ^#^P<0.05 versus SF1-Cre/NaChBac. **(c)** Time spent in the light room. N =8-10 per group; one-way ANOVA; **P<0.01, ***P<0.001 versus EYFP. **(d)** Nesting scores over time. n = 6-7 per group; two-way ANOVA; *P<0.05, **P<0.01 versus EYFP at 4 h. **(e)** Time spent interacting with a stranger mouse. n = 7-10 per group; one-way ANOVA; ***P<0.001 versus EYFP.

**Figure S3.**
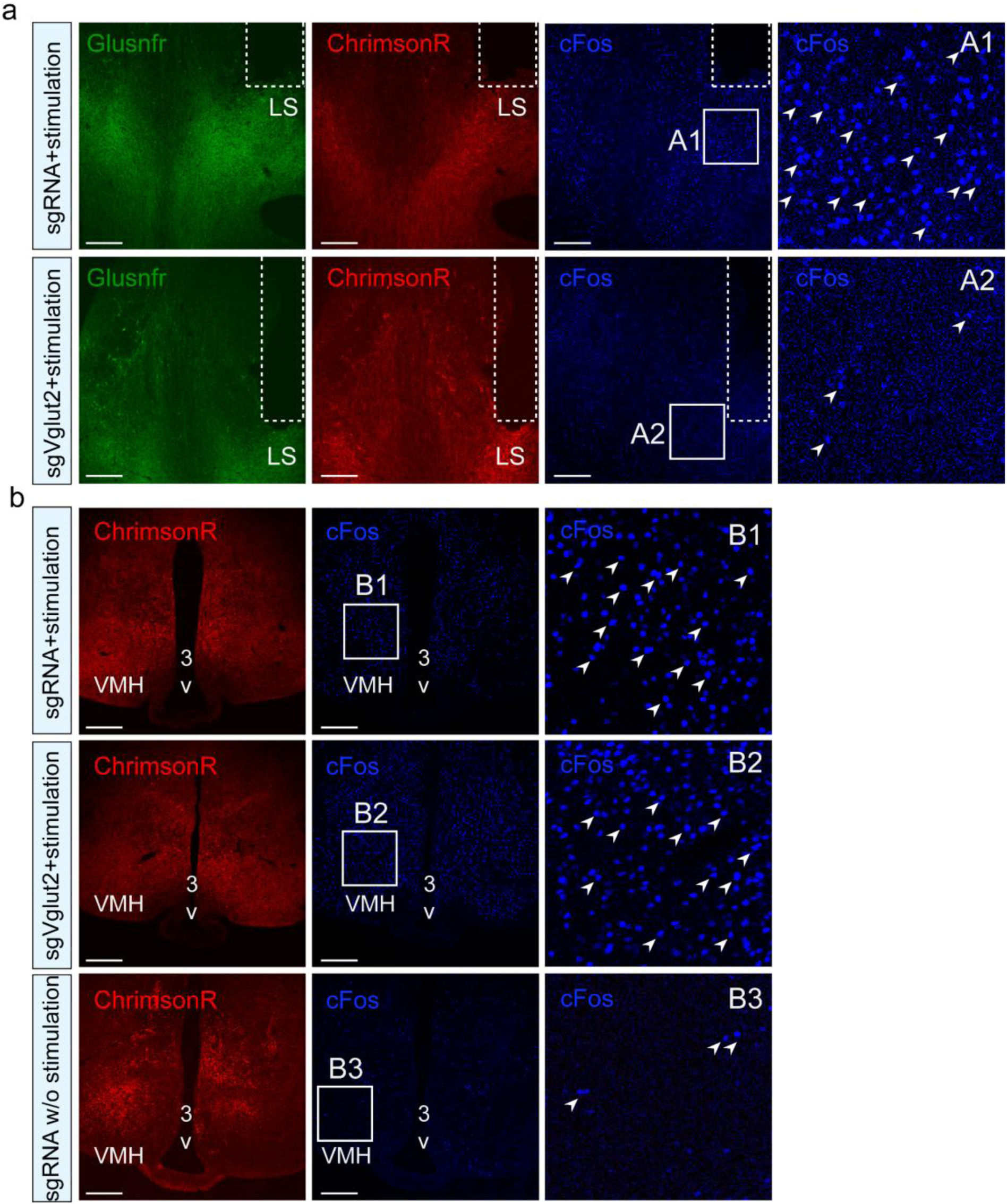
cFos immunostaining following optogenetic stimulation of VMH neuronal terminals in the LS (related to Fig.3). Mice received optogenetic photostimulation of VMH neuronal terminals in the LS (630 nm, 10 Hz, 50 ms, 30 mW, 10 min) or no stimulation. One hour after stimulation, mice were sacrificed for cFos immunostaining. **(a)** Representative images of cFos expression in the LS following photostimulation. **(b)** Representative images of cFos expression in the VMH in mice with or without photostimulation. Scale bar, 250 μm.

**Figure S4.**
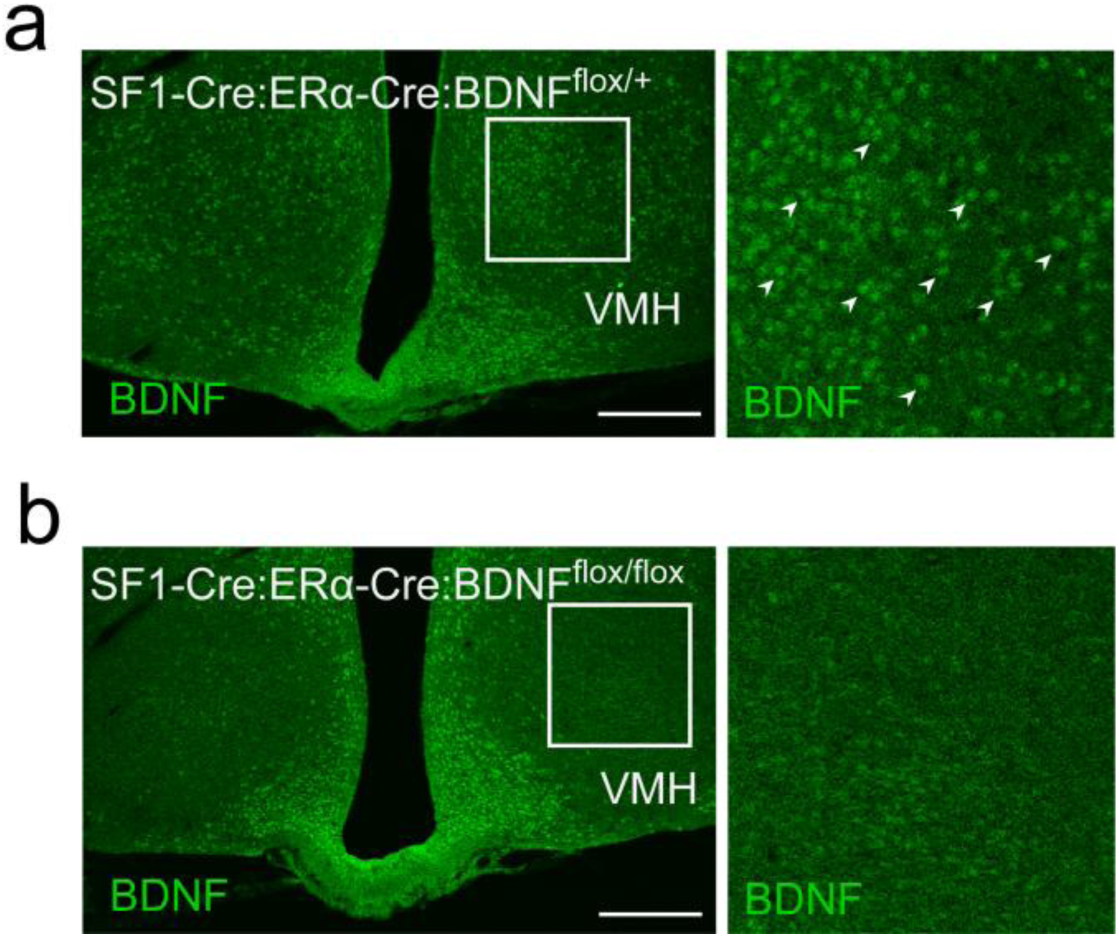
Validation of BDNF knockout (related to Fig.4). BDNF immunostaining in SF1-Cre::ERa-Cre::BDNF^flox/+^ **(a)** and SF1-Cre::ERa-Cre::BDNF^flox/flox^ **(b)** mice in VMH region. Arrows indicate BDNF positive cells. Scale bar=250 μm.

**Figure S5.**
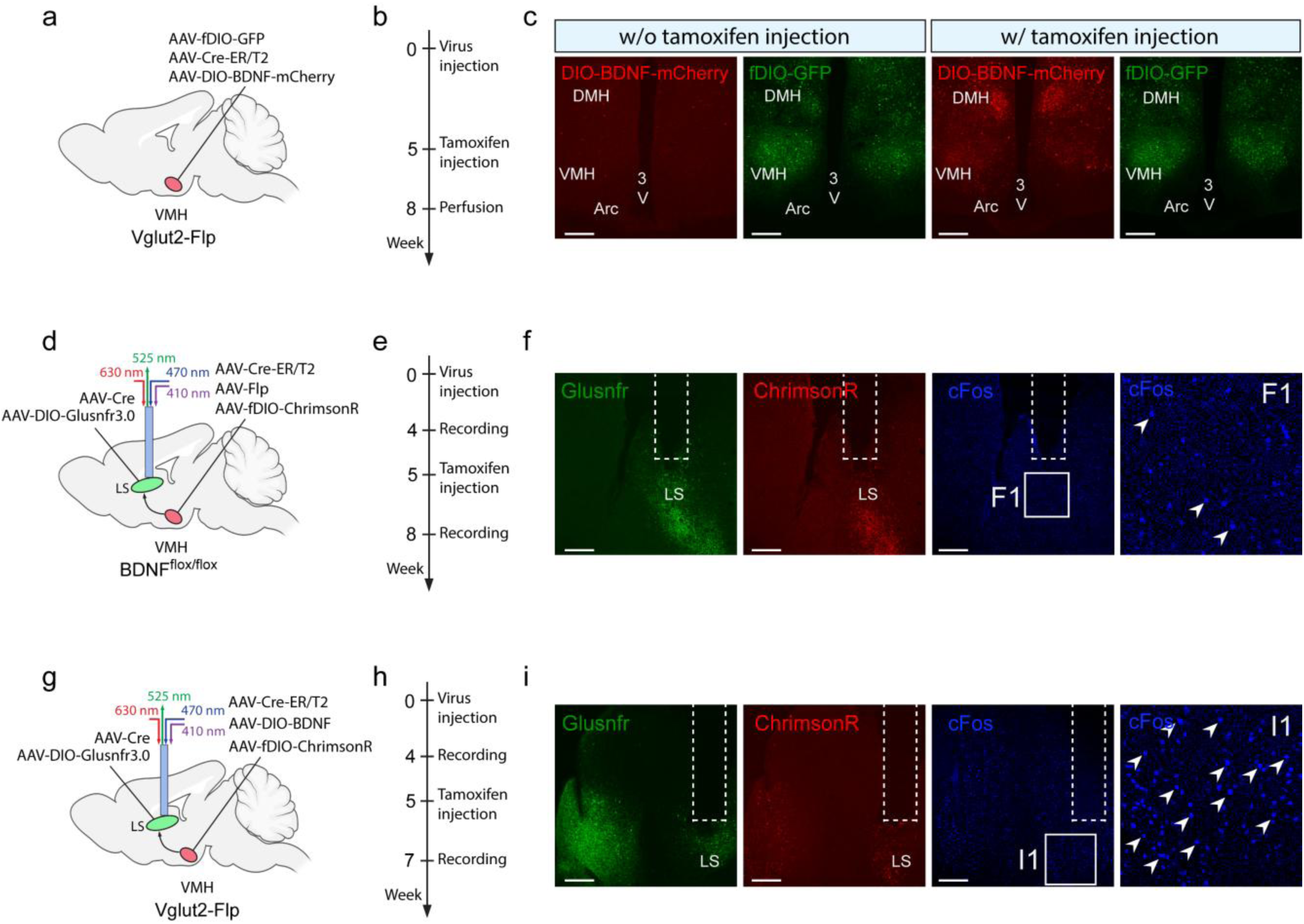
Validation of the inducible Cre-ER/T2 system (related to Fig.4 and Fig. 6). **(a)** Schematic of AAV-fDIO-GFP, AAV-Cre-ER/T2, and AAV-DIO-BDNF-mCherry injections into the VMH of Vglut2-Flp mice. **(b)** Experimental timeline showing tamoxifen injection. **(c)** Representative expression of DIO-BDNF-mCherry and fDIO-GFP in mice with or without tamoxifen treatment. **(d–i)** Mice received optogenetic photostimulation of VMH neuronal terminals in the LS (630 nm, 10 Hz, 50 ms, 30 mW, 10 min). One hour after photostimulation, mice were sacrificed for cFos immunostaining. **(d)** Schematic of inducible BDNF deletion in BDNF^flox/flox^ mice. **(e)** Experimental timeline showing tamoxifen injection. **(f)** Representative cFos expression in the LS following photostimulation in mice with inducible BDNF deletion. **(g)** Schematic of inducible BDNF overexpression in Vglut2-Flp mice. **(h)** Experimental timeline showing tamoxifen injection. **(i)** Representative cFos expression in the LS following photostimulation in mice with inducible BDNF overexpression. Scale bar, 250 μm.

**Figure S6.**
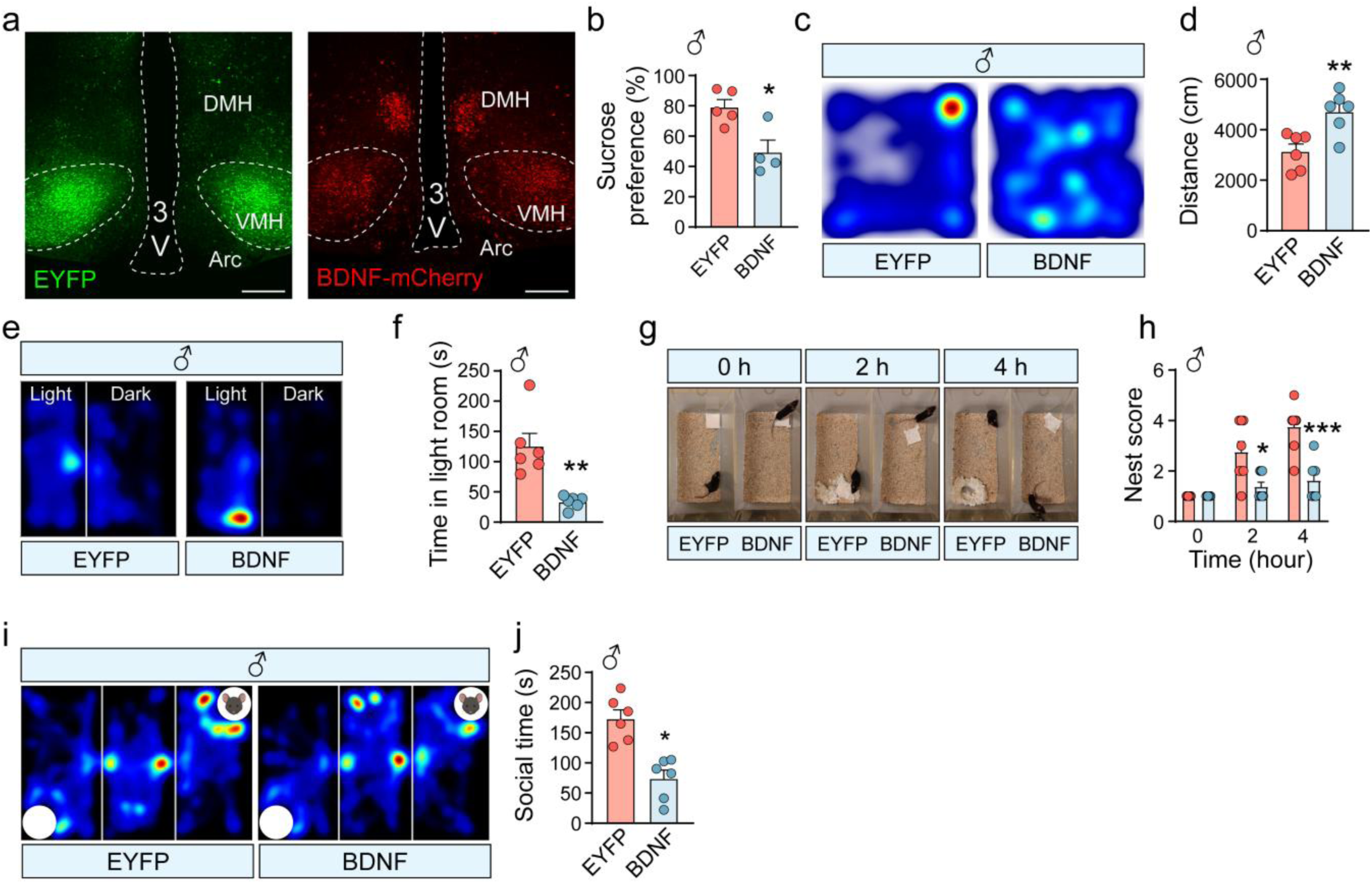
Augmented BDNF action from VH glutamatergic neurons in males caused anorexia phenotypes (related to Fig. 5). **(a)** Representative images showing EYFP (green) and BDNF-mCherry (red) expessing pattern in VH region of Vglut2-Cre male mice. Scale bar=200 μm. **(b)** Sucrose preference in EYFP- and BDNF-expressing male mice. n=5 (EYFP), n=4 (BDNF); unpaired two-tailed t test, P=0.013. **(c, d)** Open-field test. **(c)** Representative movement trajectories. **(d)** Total distance traveled in 15 min. n = 6 per group; unpaired two-tailed t test, P=0.006. **(e, f)** Light-dark box test. **(e)** Representative movement trajectories. **(f)** Time spent in the light room. n =6 per group; unpaired two-tailed t test, P=0.002. **(g, h)** Nesting behavior. **(g)** Representative images. **(h)** Nesting scores over time. n = 6 per group; two-way ANOVA; P=0.038 at 2 h, P<0.001 at 4 h. **(i, j)** Three-chamber sociability test. **(i)** Representative trajectories. **(j)** Time spent interacting with a stranger mouse. n = 6 per group; unpaired two-tailed t test; P<0.001.

**Figure S7.**
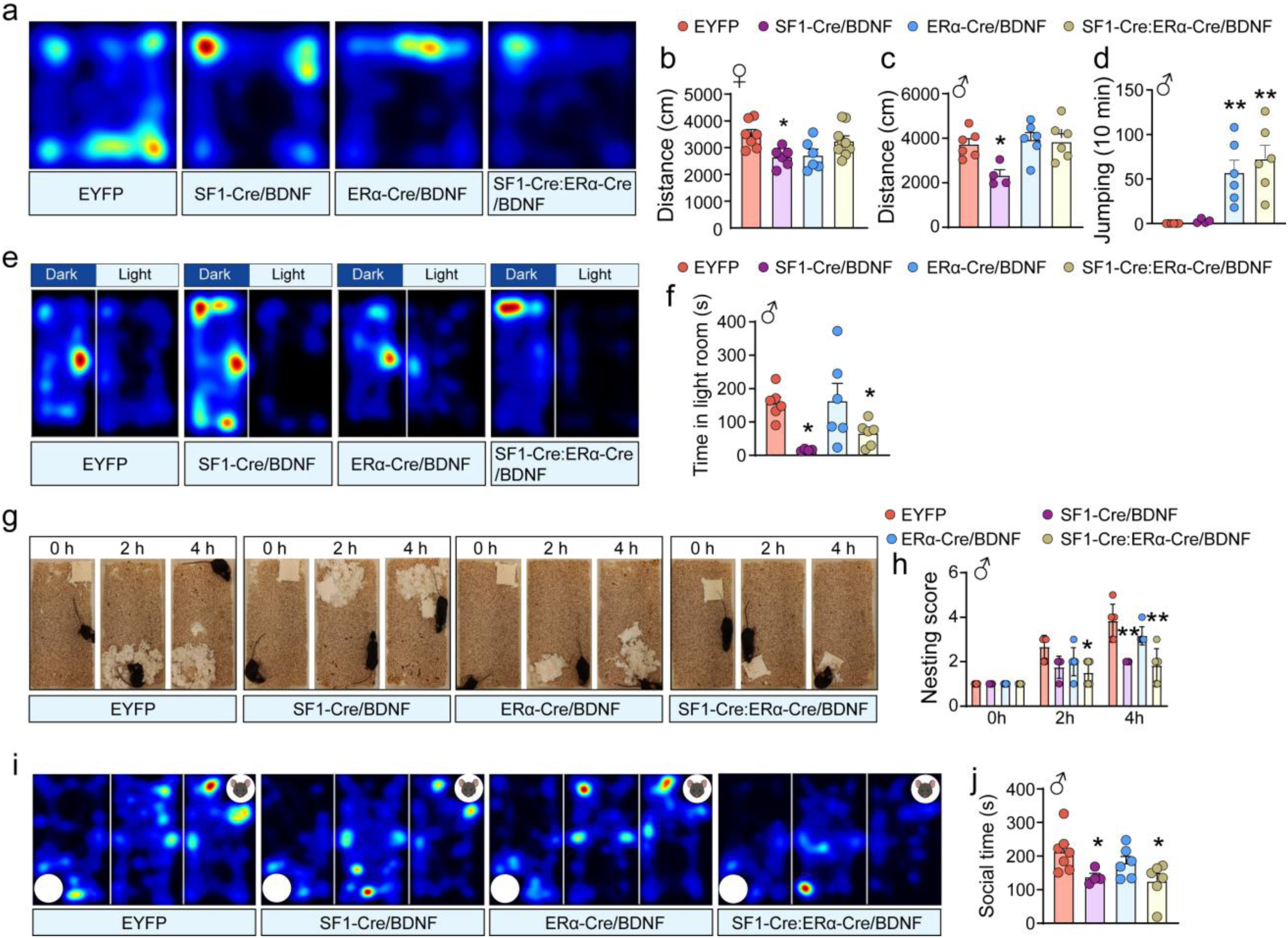
Augmented BDNF action from SF1/ERa neurons in males led to anorexia phenotypes (related to Fig. 6). **(a-c)** Open-field test. **(a)** Representative movement trajectories. Total distance traveled in 15 min in female **(b)** and male **(c)** mice. Females, n=6-8 per group; males, n=4-6 per group; one-way ANOVA; * P<0.05 versus EYFP. **(d)** Total number of jumps during the 10-min test in male mice. n=6 (EYFP), n=4 (SF1-Cre+BDNF), n=6 (ERa-Cre+BDNF), n=6 (SF1-Cre::ERa-Cre+BDNF); one-way ANOVA; **P<0.01 versus EYFP. **(e, f)** Light-dark box test. **(e)** Representative movement trajectories. **(f)** Time spent in the light room. n =4-6 per group; one-way ANOVA, *P<0.05 versus EYFP. **(g, h)** Nesting behavior. **(g)** Representative images. **(h)** Nesting scores over time. n = 4-6 per group; two-way ANOVA; *P<0.05, **P<0.01 versus EYFP. **(i, j)** Three-chamber sociability test in male mice. **(i)** Representative trajectories. **(j)** Time spent interacting with a stranger mouse. n = 4-6 per group; *P<0.05 versus EYFP.

**Figure S8.**
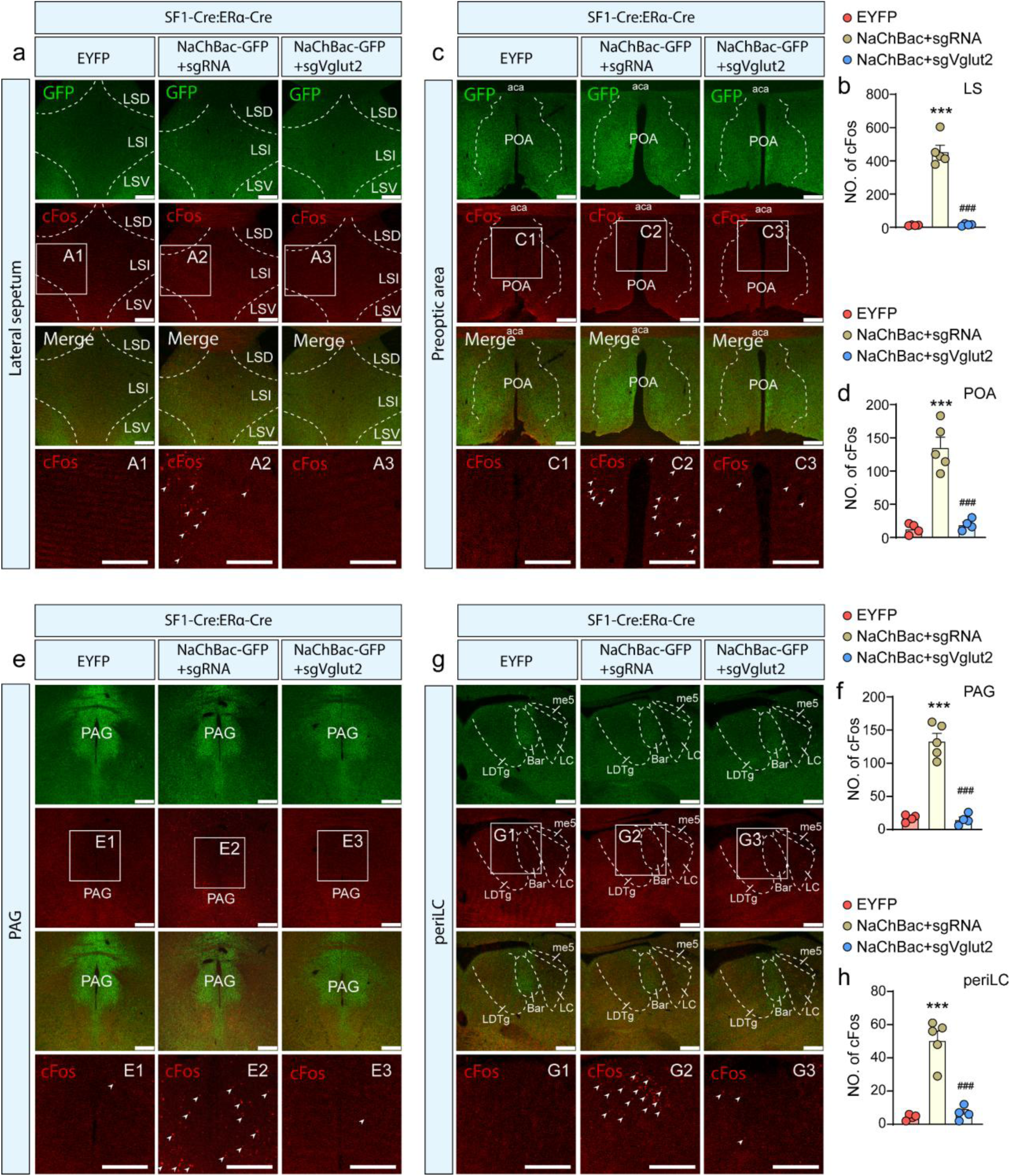
c-Fos in downstream neurons of VH in AAV-DIO-EYFP, AAV-DIO-NaChBac+sgRNA, AAV-DIO-NaChBac+sgVglut2 injection SF1-Cre::ERa-Cre mice (related to Fig.7). Representative images showing GFP (green) and c-Fos (red, arrows) expression pattern in lateral septum **(a)**, preoptic area **(c)**, PAG **(e)**, periLC **(g)**. scale bar=250 μm. The pictures in the bottom row of each panel represent the magnified pictures shown in the boxed

**Figure S9.**
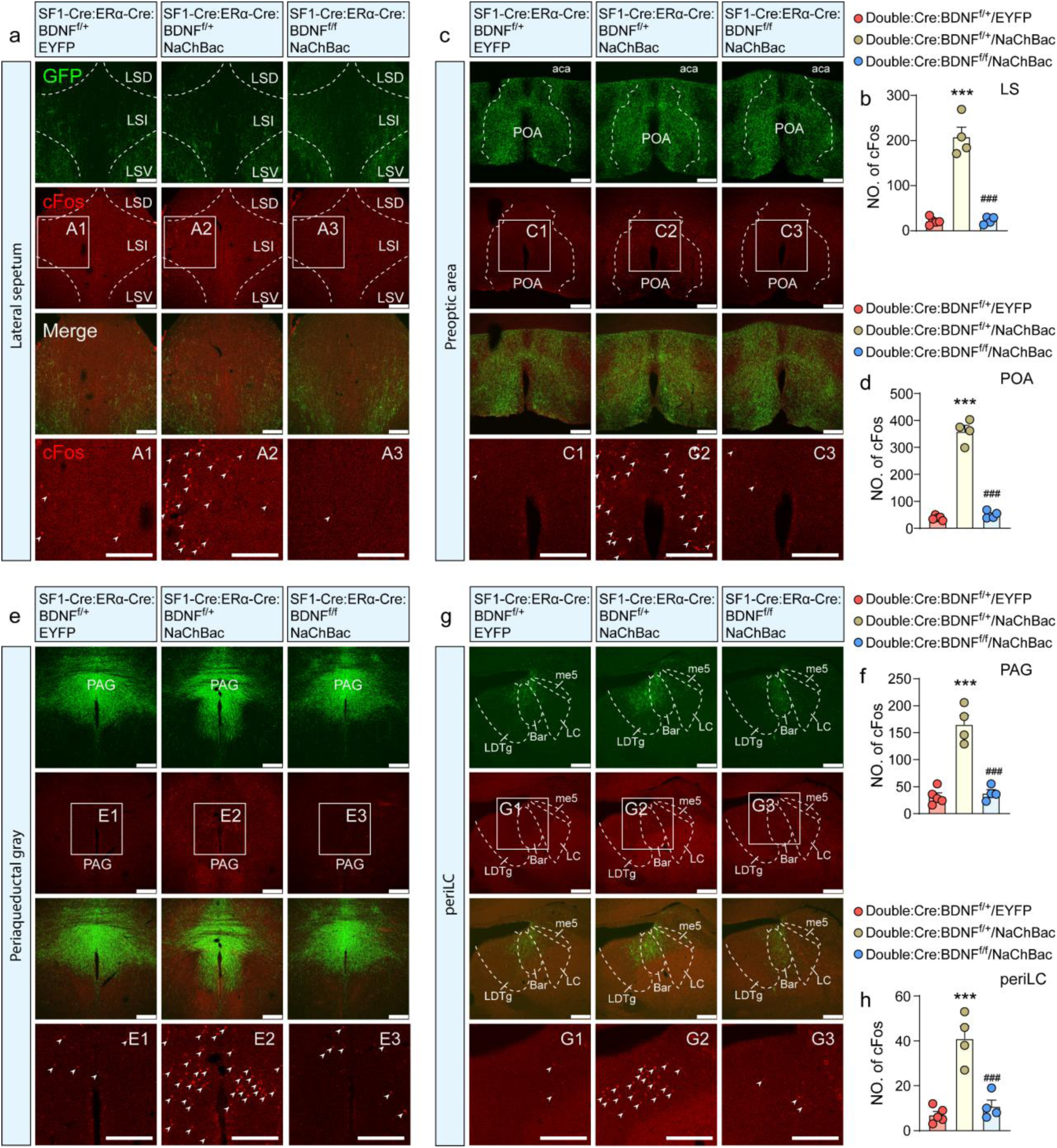
c-Fos in downstream neurons of VH in EYFP/SF1-Cre::ERa-Cre::BDNFflox/+, NaChBac/SF1-Cre::ERa-Cre::BDNF^flox/+^, and NaChBac/SF1-Cre::ERa-Cre::BDNF^flox/flox^ mice (related to Fig.6). Representative images showing GFP (green) and c-Fos (red, arrows) expression patterns in lateral septum **(a)**, preoptic area **(c)**, PAG **(e)**, periLC **(g)**. scale bar=250 μm. The pictures in the bottom row of each panel represent the magnified pictures shown in the boxed

**Figure S10.**
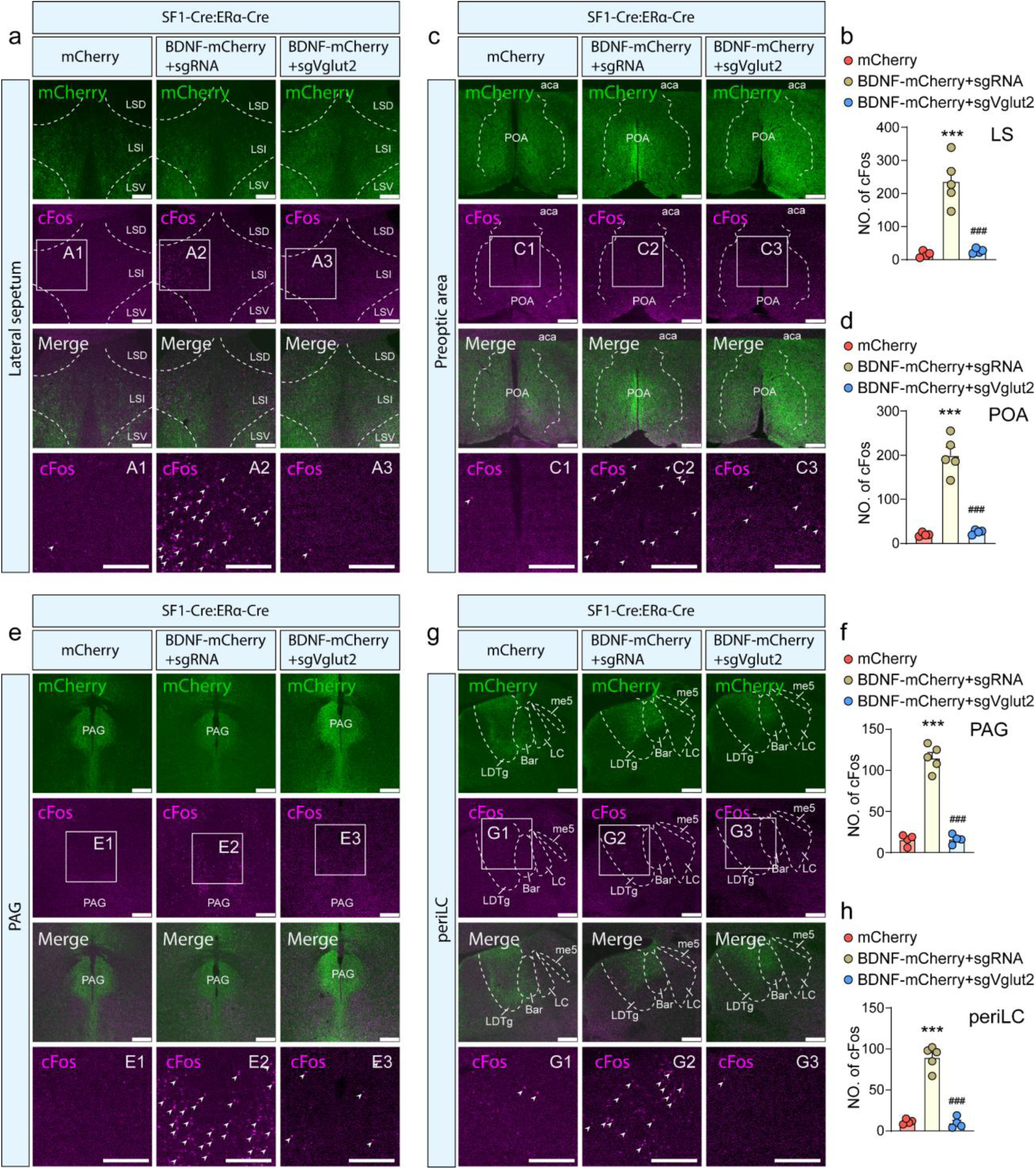
c-Fos in downstream neurons of VH in mCherry, BDNF+sgRNA, BDNF+sgVglut2 virus injected SF1-Cre::ERa-Cre mice (related to Fig.7). Representative images showing mCherry (green) and c-Fos (magenta, arrows) expression pattern in lateral septum **(a)**, preoptic area **(c)**, PAG **(e)**, periLC **(g)**. scale bar=250 μm. The pictures in the bottom row of each panel represent the magnified pictures shown in the boxed area in each column. Number of cFos positive cells in in lateral septum **(b)**, preoptic area **(d)**, PAG **(f)**, periLC **(h)**. n=4-5 per group; one-way ANOVA; ***P<0.001 versus mCherry; ^###^P<0.001 versus BDNF+sgRNA.

**Figure S11.**
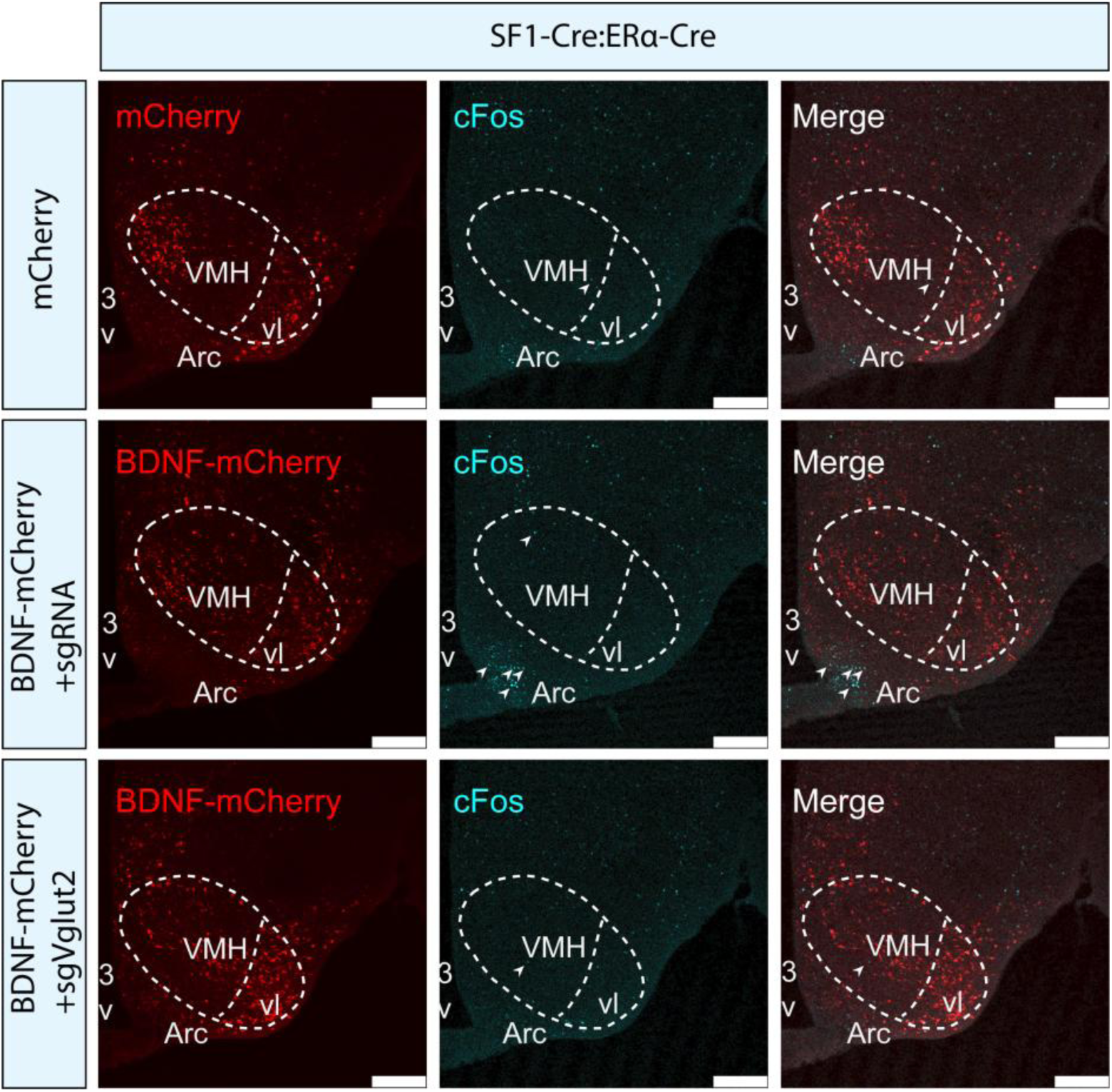
Representative images showing mCherry (red) and cFos (cyan, arrows) expression pattern in VH in mCherry, BDNF+sgRNA, BDNF+sgVglut2 injection SF1-Cre::ERa-Cre mice (related to Fig.7). Scale bar=200 μm.

**Figure S12.**
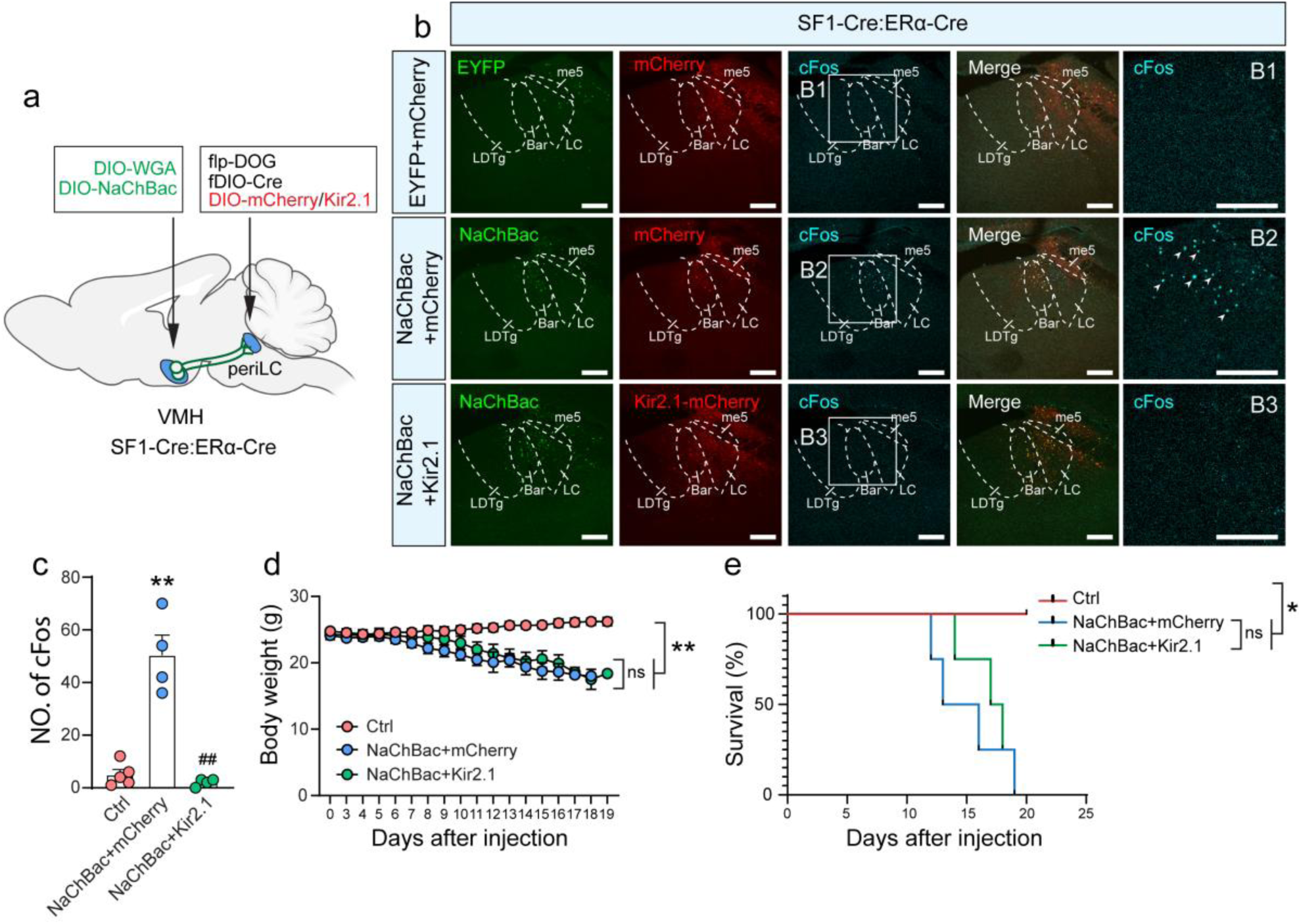
Inhibition of neurons in periLC, a downstream of VH, doesn’t improve AN phenotypes (related to Fig.7). **(a)** Diagram of viral delivery (AAV-DIO-WGA, AAV-DIO-NaChBac) into VH, and flp-DOG, fDIO-Cre, DIO-mCherry/Kir2.1 into periLC in SF1-Cre::ERa-Cre mice. **(b)** Representative images showing GFP (green), mCherry (red), Kir2.1 (red), and cFos (cyan, arrows) expression pattern in periLC. Scale bar=250 μm. The pictures in the right column represent the magnified pictures shown in the boxed area in the corresponding box in the same row. **(c)** Number of c-Fos positive cells in periLC. n=4-5 per group; one-way ANOVA; **P<0.01 versus Ctrl; ^##^P<0.01 versus NaChBac+mCherry. **(d)** Body weight trajectories of mice. n=4-5 per group, two-way ANOVA, **P<0.01. **(e)** Kaplan–Meier survival analysis of mice. Mice were considered dead when body weight loss reached 30%, log-rank test, *P<0.05 versus EYFP.

